# Spatiotemporal patterns of urban mosquitoes are modulated by socioeconomic status and environmental traits in the United States

**DOI:** 10.1101/2021.10.30.466623

**Authors:** Senay Yitbarek, Kelvin Chen, Modeline Celestin, Matthew McCary

## Abstract

The distribution of mosquitoes and associated vector diseases (e.g., West Nile, dengue, and Zika viruses) is likely a function of environmental conditions in the landscape. Urban environments are highly heterogeneous in the amount of vegetation, standing water, and concrete structures covering the land at a given time, each having the capacity to influence mosquito abundance and disease transmission. Previous research suggests that socioeconomic status is correlated with the ecology of the landscape, with lower-income neighborhoods generally having more concrete structures and standing water via residential abandonment, garbage dumps, and inadequate sewage. Whether these socio-ecological factors affect mosquito distributions across urban environments in the United States (US) remains unclear. Here, we present a meta-analysis of 22 paired observations from 15 articles testing how socioeconomic status relates to overall mosquito burden in urban landscapes in the United States. We then analyzed a comprehensive dataset from a socioeconomic gradient in Baltimore, Maryland to model spatiotemporal patterns of *Aedes albopictus* using a spatial regression model with socio-ecological covariates. The meta-analysis revealed that lower-income neighborhoods (regions making less than $50,000 per year on average) are exposed to 151% greater mosquito densities and mosquito-borne illnesses compared to higher-income neighborhoods (≥$50,000 per year). Two species of mosquito (*Ae. albopictus* and *Aedes aegypti*) showed the strongest relationship with socioeconomic status, with *Ae. albopictus* and *Ae. aegypti* being 62% and 22% higher in low-income neighborhoods, respectively. In the spatial regression analysis in Baltimore, we found that *Ae. albopictus* spatial spread of 1.2 km per year was significantly associated with median household income, vegetation cover, tree density, and abandoned buildings. Specifically, *Ae. albopictus* abundance was negatively correlated with median household income, vegetation cover, and tree density. *Ae. albopictus* abundance and the cover of abandoned buildings were positively correlated. Together, these results indicate that socio-ecological interactions can lead to disproportionate impacts of mosquitoes on humans in urban landscapes. Thus, concerted efforts to manage mosquito populations in low-income urban neighborhoods are required to reduce mosquito burden for the communities most vulnerable to human disease.

## INTRODUCTION

Mosquitoes (Diptera: Culicidae) are a significant threat to humans, killing an estimated 780,000 people per year worldwide (WHO, 2014). Mosquito burden, the harm mosquitoes inflict on humans through bites and vectored diseases, is expected to intensify as global change progresses (Whiteman et al. 2020, Holeva-Eklund et al. 2021). Since mosquitoes are responsible for 36% of all vector-borne diseases globally (Swei et al. 2020), managing their impacts is imperative for global health (Leisnham and Juliano 2012, Mavian et al. 2019a).

A primary driver of global change is urban expansion resulting from human population growth, with projections indicating 70% of the world population living in cities by 2050 (Heilig 2012, Huang et al. 2019). Urban environments also harbor invasive, medically important mosquitoes, such as the mosquito genus *Aedes*, which are competent disease vectors of chikungunya, dengue, yellow fever, and Zika (Goodman et al. 2018, Rose et al. 2020). Because urban environments are highly heterogeneous due to social and ecological factors (Marzluff 2008, McKinney 2008), mosquitoes can be unevenly distributed within cities (Little et al. 2017). Mosquito hotspots generally emerge with favorable conditions: neighborhoods with abundant breeding habitats, high densities of human hosts, and low mosquito-mitigation efforts (LaDeau et al. 2013a, Faraji et al. 2014). While developed countries have successfully eliminated some pathogens such as malaria (Benelli 2015), mosquito-borne diseases remain a public health threat in temperate regions, such as the United States (US), where cities can be large heat islands supporting high densities of mosquitoes and vector diseases (Mavian et al. 2019b).

Much of what is known about mosquito effects in cities relate to how environmental factors influence human exposure to mosquitoes (reviewed in Sallam 2017, Whitney 2020, Whiteman 2020). In urban environments, mosquito populations are affected by the availability of water-holding containers, vegetation, and human population density—particularly for the mosquitoes *Aedes. albopictus*, *Ae. aegypti*, and *Culex* spp. More recently, however, studies have investigated whether these ecological attributes in cities relate to socioeconomic status (SES), with mixed results (Sallam 2017, Whiteman 2020). For example, some studies report higher mosquito densities in low SES neighborhoods in cities (LaDeau et al. 2013b, Dowling et al. 2013, Little et al. 2017), whereas other studies show no relationship between low and high SES communities (Van de Poel et al. 2009, Whiteman et al. 2020). Such inconclusive findings make it difficult to generalize mosquito burden across urban environments, limiting our ability to predict their impacts as urban expansion advances. One way to address the contradictory findings is to perform a quantitative synthesis to understand their net effects regarding how urban mosquitoes associate with SES (Koricheva and Gurevitch 2013).

While meta-analytic methods are useful for describing patterns across studies (Koricheva et al. 2014), they are typically crude and do not yield mechanistic insights. Furthermore, with respect to urban mosquitoes, the environmental covariates underpinning mosquito spatiotemporal patterns and spread are wide-ranging and remain largely unexplored (Sallam et al. 2017). *Ae. albopictus*, the invasive tiger mosquito, is a widespread urban mosquito that was established in the US in the 1980s (Benedict et al. 2007). The Invasive Species Specialist Group considers it a top 100 invasive species, and it is now the most invasive mosquito species worldwide (Kotsakiozi et al. 2017, Cuthbert et al. 2021). Previous research links a handful of environmental covariates to *Ae. albopictus* populations in urban environments (Little et al. 2017), suggesting that *Ae. albopictus* populations are more affected by building abandonment and vegetation and that those factors can interact with SES. Whether these environmental covariates can be used to predict *Ae. albopictus* spatiotemporal patterns and spread across urban environments remains unknown. The speed and spread of *Ae. albopictus* into new environments will depend on the combined and interactive effects of dispersal, population growth, and availability of suitable habitats. Although previous models (e.g., species distribution models) have successfully estimated the timing and spatial spread of invasive species (Guisan and Thuiller 2005, Elith and Leathwick 2009), they do not fully account for local spread rates concerning environmental characteristics.

In the present study, we employ the first meta-analysis to investigate how SES in US-based urban environments correlates with mosquito densities and vector-disease transmission. We then used a comprehensive dataset from an SES gradient in Baltimore, Maryland (Little et al. 2017) to model the distribution and spread of the invasive mosquito *Ae. albopictus* using a spatial regression model. We hypothesized that if SES encourages uneven mosquito burden in urban environments because they correlate with environmental conditions more suitable to mosquito populations (Fig. 1), we expect to find higher mosquito burden in low SES than high SES neighborhoods. Furthermore, we suspect the environmental covariates that predict *Ae. albopictus* spread and speed will be linked indirectly to SES (Fig. 1). The overall goal of this research is to understand how SES and ecological factors relate to mosquito populations across local and regional scales.

**Figure 1.**
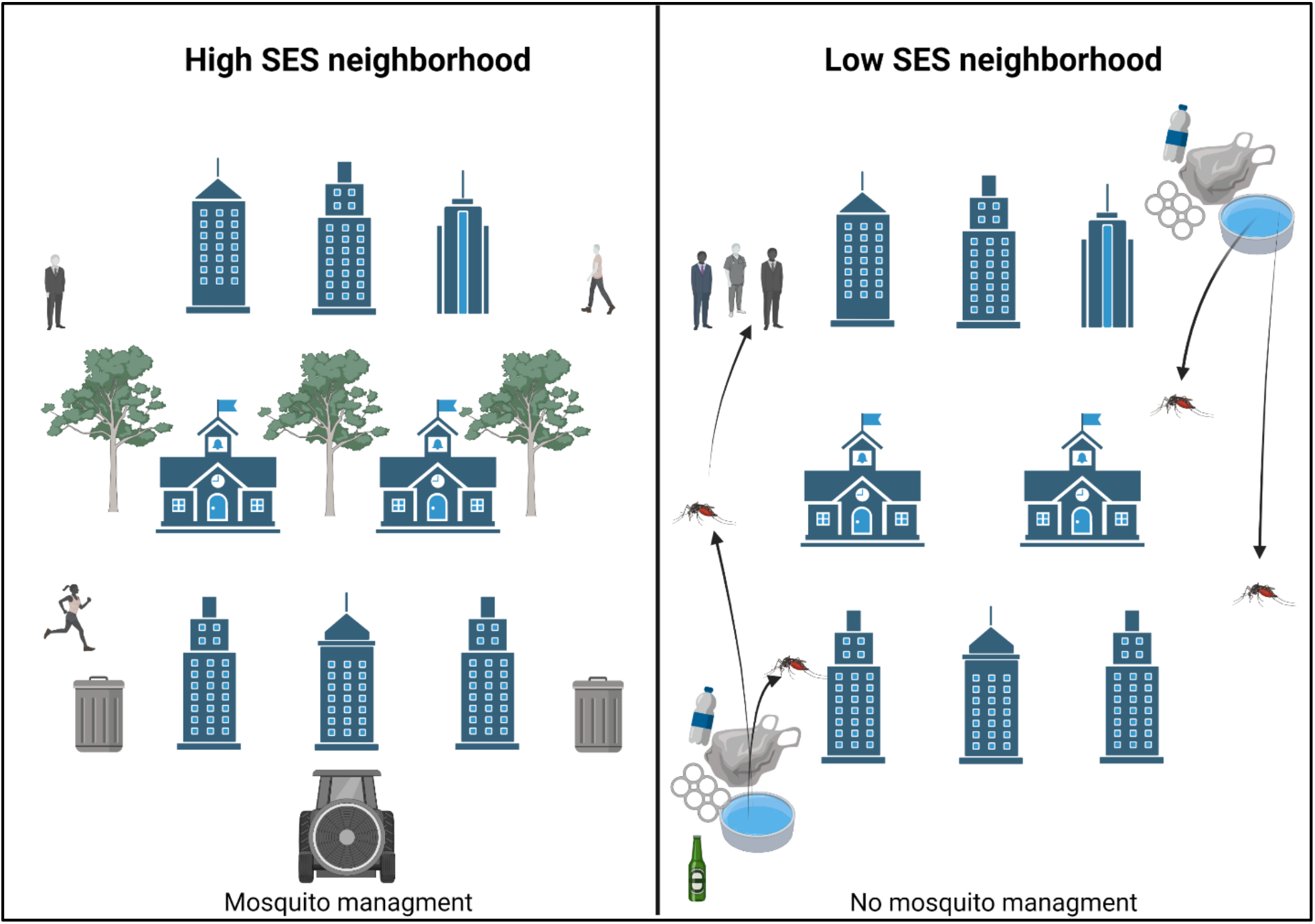
Conceptual diagram illustrating how mosquito burden can be mediated by socioeconomic status (SES) and environmental traits. In low SES neighborhoods, the presence of garbage, water-holding containers, poor sewage, and absence of mosquito-control programs allow for urban mosquitoes to proliferate in high numbers. In contrast, high SES neighborhoods are highly managed for mosquitoes and often have fewer artificial structures that can encourage mosquito populations.

## METHODS

### Study 1: Meta-analysis of mosquito burden in US-based urban environments

#### Literature search and data extraction

To uncover the studies that evaluated how SES in urban environments correlated with mosquito densities and vector-disease transmission in humans, we conducted a literature search for primary articles in ISI Web of Science. We used the following string of search terms without any restriction on the year of publication (last accessed on 13 August 2021): (Socioeconomic OR socio economic OR socio-economic OR socio ecological OR socio-ecological OR wealth OR income OR poverty) AND (urban* OR cit* OR town* OR population* OR densit* OR minorit* OR communit* OR neighborhood*) AND (vector* OR virus* OR West Nile OR Dengue OR chikungunya OR malaria OR Zika OR disease*) AND (Aedes aegypti OR Aedes albopictus OR Culex tarsalis OR Culex quinquefasciatus OR Anopheles freeborni OR Anopheles quadrimaculatus OR mosquito*) AND (United States OR USA* OR North America*). Our initial search resulted in 322 published articles. We reviewed each article’s titles and abstracts to discern its potential fit for inclusion in the meta-analysis. Next, we assessed each eligible article’s reference lists to uncover other potentially pertinent articles and contacted a handful of experts in the US regarding available research articles. We used no unpublished data sets in this meta-analysis. Although we acknowledge that we might not have collected every possible article, these search methods provided adequate coverage of the primary literature. Lastly, we recorded each article that emerged from the literature search and documented the number of studies excluded based on our inclusion criteria (Appendix S1: Tables S1-S2). Our meta-analysis search protocol followed the Preferred Reporting Items for Systematic Reviews and Meta-analysis (PRISMA; Moher et al. 2009) (Fig. 2).

**Figure 2.**
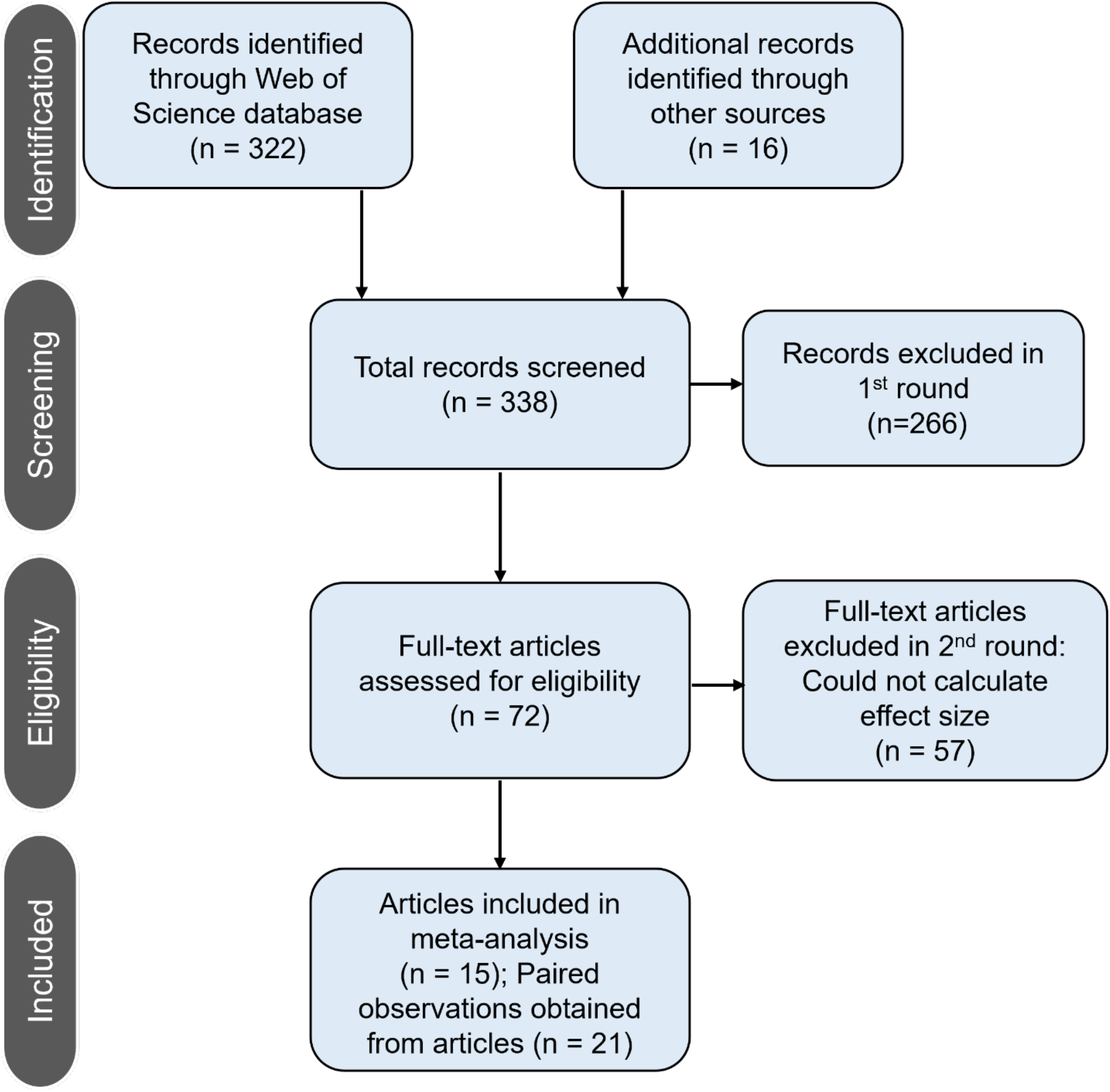
The modified PRISMA flow schematic (Moher et al. 2009). We recorded the number of articles screen during each step of the literature survey and meta-analysis

The meta-analysis was designed to evaluate peer-reviewed studies that compared mosquito densities or mosquito-borne diseases in low SES and high SES urban neighborhoods in the US (Table 1). Here, we defined SES using the information provided in the original study. If the study did not define SES for the urban environments, we characterized low SES as households in neighborhoods making less than $50,000 per year and high SES as neighborhoods making ≥$50,000 per year (inflation was accounted for in older studies). Nevertheless, almost all the eligible studies fell within these two SES categories. Finally, we only used articles in which the original study explicitly stated it investigated a city, metropolitan, and/or urban environment.

**Table 1.** The characteristics of the original studies included in the meta-analysis.

| Study | Year | City/Region | State | SES<br>Low | SES<br>High | Taxon | Development<br>stage | Data type |
| --- | --- | --- | --- | --- | --- | --- | --- | --- |
| (1) Barrera et al. | 2021 | Southern<br>Puerto Rico | Puerto Rico | NA | NA | Ae.<br>aegypti,<br>Culex spp. | Pupae | Abundance |
| (2) Becker et al. | 2014 | Baltimore | Maryland | NA | NA | Culex spp.,<br>Ae.<br>albopictus | Adult | Abundance |
| (3) Chambers et al. | 1986 | East Baton<br>Rouge | Louisiana | NA | NA | Ae. aegypti | Larvae | Frequency |
| (4) Crespo et al. 2021 | 2021 | Baton Rouge | Louisiana | <35<br>K | <35K | Culex spp.,<br>Ae.<br>albopictus,<br>Anopheles<br>spp. | Adult | Abundance |
| (5) Donnelly et al. | 2020 | Los Angeles | California | 42K | 75K | Ae. aegypti | Adult | Abundance |
| (6) Goodman et al. | 2018 | Baltimore | Maryland | <50<br>K | >50K | Ae.<br>albopictus | Adult | Blood meal<br>frequency |
| (7) LaDeau et al. | 2013 | Washington,<br>DC | Washington,<br>DC | <50<br>K | >50K | Mosquitoes<br>indist. | Pupae | Frequency |
| (8) Little et al. | 2017 | Baltimore | Maryland | <50<br>K | >50K | Ae.<br>albopictus | Larvae | Frequency |
| (9) Little et al. | 2021 | Suffolk | Virginia | <40<br>K | >57K | Ae.<br>albopictus | Adult | Blood meal<br>frequency |
| (10) Martin et al. | 2019 | Lower Rio Grande Valley | Texas | <50 K | >50K | Ae. aegypti | Adult | Abundance |
| (11) Ruiz et al. | 2007 | Chicago | Illinois | <50 K | >50K | Mosquitoes indist. | Adult | Frequency |
| (12) Scavo et al. | 2021 | San Juan | Puerto Rico | <35 K | >50K | Mosquitoes indist. | Adult | Diversity index |
| (13) Shragai and Harrington | 2019 | Southern New York | New York | <50 K | >110K | Ae. albopictus | Adult | Frequency |
| (14) Unlu et al. | 2011 | Mercer, Monmouth County | New Jersey | <50 K | >50K | Ae. albopictus | Adult | Abundance |
| (15) Walker et al. | 2018 | Tucson | Arizona | <35 K | >50K | Ae. aegypti | Larvae | Frequency |

We focused the meta-analysis on the US because it: (1) allows for a more objective comparison of low and high SES neighborhoods, given that categories of SES can vary widely across countries, and (2) makes it easier to relate the results of the meta-analysis to the results from the spatial modeling of urban mosquitoes in Baltimore, Maryland (see Study 2 below). Each study had to compare mosquito burden in low vs. high (or medium) SES neighborhoods. Here, mosquito burden could be the number of mosquito larvae, adults, confirmed diagnosis of mosquito-borne illnesses (e.g., West Nile, dengue, and Zika viruses), and proportion of mosquito human bloodmeals in the population.

The articles eligible for our meta-analysis required the following criteria: (1) A comparison between high and low SES urban neighborhoods. When multiple units of high and low SES neighborhoods were provided, we calculated a composite average of mosquito burden for each mosquito species across SES. (2) If a study provided multiple sample dates for mosquito burden, we only used the final period. (3) The studies had to include at least the mean and sample size for both high and low SES neighborhoods. For the studies that did not include an estimate of variance, we employed a linear regression model using the studies with complete information to fill in missing variance values in the other articles (i.e., an imputation technique; Koricheva et al. 2013). Our regression model proved to be a good predictor for the missing variance values (*R*^2^ = 0.65, *p* < 0.001). (4) Each study needed to be an original research article. We did not include other meta-analyses, reviews, or modeling papers. Also, we only used one article if multiple publications used the same dataset.

Data points were obtained from the text, tables, supplemental materials, and figures. Data from figures were extracted with ImageJ (Abràmoff et al. 2004), an image processing software.

#### Analysis

We used the log response ratio (LRR) (Hedges et al. 1999) effect size to measure the influence of SES on mosquito burden by calculating:

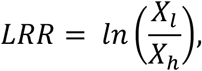

where X_l_ and X_h_ are the sample means of low and high SES neighborhoods, respectively. A positive LRR denotes that mosquito burden in low SES neighborhoods is higher, whereas a negative LRR means that high SES neighborhoods experienced higher mosquito burden. The LRR variance was calculated as:

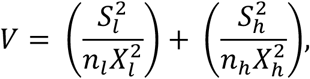

where S and n denote the standard deviation and sample size of replicates, respectively. The subscript ‘l’ and ‘h’ refer to the low and high SES neighborhoods, respectively. LRR is a widely used effect size measure that allows comparisons of studies with different techniques and data types (Hedges et al. 1999, Lajeunesse 2015).

We conducted a random-effects model (REM) of meta-analysis to determine the overall effect of SES on mosquito burden in urban environments (Borenstein et al. 2011). Next, we performed a mixed-effects model (MEM) with restricted maximum likelihood (REML) to evaluate differences in the effects of SES with mosquito taxa as moderators and using weighted mean effect sizes (Borenstein et al. 2011). Here, the heterogeneity of effect sizes was calculated through the Q statistic, which is used to estimate the amount of heterogeneity attributed to unexplained variation due to unknown differences in environmental conditions across the studies (i.e., the weighted sums of squares tested against a χ2 distribution; Hedges and Olkin 1985). We considered mean effect sizes as statistically different if their 95% confidence intervals (CI) did not include zero (Borenstein et al. 2011). We used the ‘metafor’ package in R version 4.0.3 (R Development Core Team 2020) to conduct the meta-analysis.

#### Publication bias

Because studies with significant results are more likely to get published, the primary literature on a given subject can underrepresent studies reporting non-significant results, leading to publication bias (Jennions et al. 2013). To test if our results were affected by publication bias, we employed two methods. First, we used Trim-and-Fill funnel plots (Duval and Tweedie 2000), which are plots that illustrate effect sizes against sample sizes from individual studies. Trim-and-Fill plots from our data show symmetrical scatter plots across the 22 paired observations, indicating no evidence of publication bias (Appendix S2: Fig. S1). Second, we calculated Rosenthal’s fail-safe number (Orwin 1983) for our random-effects models. Rosenthal’s fail-safe number is the number of missing case studies with non-significant results needed to nullify the combined effect size (Orwin 1983). Our results also suggest no evidence of publication bias using Rosenthal’s fail-safe number (number of studies to nullify the result is 441; *p* < 0.001, Appendix S1: Table S1)

### Study 2: Spatiotemporal patterns and spread of urban mosquitoes in Baltimore

#### Baltimore dataset and sample collection

We used a comprehensive dataset from Baltimore, Maryland that investigated how SES and environmental traits affected mosquito abundance (see Little et al. 2017 for full details). Briefly, adult mosquitoes and larval habitats were sampled in southwest Baltimore neighborhoods using BG-sentinel and octenol lures during the summers of 2013-2015 (Little et al. 2017). Traps were placed in pairs on 12 of the 16 focal blocks spread across five neighborhoods. The neighborhoods spanned a range of SES categorizations including high, medium, and low SES neighborhoods. Neighborhood surveys were conducted in residential areas, and SES classifications were based on median household income, educational attainment, and housing quality. All neighborhoods were similar distances from forested parks and bodies of water. Surveys were conducted to determine the number of abandoned buildings and vacant lots for each focal block.

To characterize habitat and environmental factors, total precipitation data were calculated 2 weeks before the sampling of juveniles. Normalized difference vegetation index (NDVI) was used to measure vegetation in each of the 16 block clusters for the duration of the study (2013-2015). Lastly, individual container habitats were used to proximate adult mosquito densities (Bodner 2014).

#### Analysis

This study examined the spatiotemporal distribution of *Ae. albopictus* across the highly heterogeneous environment in Baltimore, Maryland. While previous research investigating *Ae. albopictus* distributions has been limited to broad spatial and temporal scales, few studies have examined their distribution at finer spatial (e.g., within city neighborhoods) or temporal (e.g., weekly temperature observations) scales. In a previous study using this Baltimore dataset (Little et al. 2017), they applied generalized linear mixed models to evaluate how socio-ecological indicators, infrastructure, and vegetation explained the distribution of *Ae. albopictus* in the urban environment. Here, we implement recently developed methods that combine dynamic equations within a hierarchical Bayesian framework (Goldstein et al. 2018) to make predictions about the spatiotemporal distribution of *Ae. albopictus* based on social and environmental factors. We re-analyzed the Baltimore dataset (Little et al. 2017) by fitting Gaussian process gradient models; the Gaussian process was fit to *γ*(*s*). We estimated the mean and covariance parameters *Θ* = (*β*_0_, *β*_1_, *β*_2_, *σ*^2^, *ϕ*, *r*^2^) using a Bayesian approach (Banerjee et al. 2008, Goldstein et al. 2018) where *Θ* is sampled from a posterior distribution using a Markov chain Monte Carlo (MCMC) algorithm. The posterior mean is estimated as 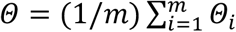. The spatial regression models using the invasion ‘Speed’ package in R (Goldstein et al. 2018) were fit to geographical coordinates of mosquito sites sampled in focal blocks across five neighborhoods. In the analysis, the response variable represents the time (2013-2015) the mosquitos were sampled; the predictor variables are geographical coordinates. The reciprocal of the response variable represents the measure of the spatial distribution of mosquitoes. The mean distribution of mosquitoes is estimated as. 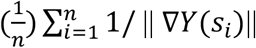 (Goldstein et al. 2021). Longitude and latitude coordinates were projected using Albers equal-area conic projection parameters. The Bayesian spatial regression model was fit using the ‘spBayes’ R package (Finley and Banerjee 2020).

## RESULTS

### Study 1: Meta-analysis of mosquito burden in US-based urban environments

Our database consisted of 15 peer-reviewed studies conducted in the US, resulting in 21 paired observations that investigated the relationship between low and high SES neighborhoods in urban areas. The studies spanned all regions of the US, including the Northeast, South, Southwest, and Midwest. Refer to Table 1 for details on the studies included in the meta-analysis.

The meta-analysis revealed that mosquito burden in low SES neighborhoods (<$50,000 USD per year on average) was much higher than high SES (≥$50,000 USD per year) neighborhoods (REM; log response ratio = 0.92, CI = [0.47, 1.36], *p* = 0.001, Fig. 3). The mixed-effects model indicated that two species of mosquito (*Ae. albopictus* and *Ae. aegypti*) showed the strongest relationship with SES (MEM, log response ratio_*Ae. albopictus*_ = 0.48, CI = [0.24, 0.72], *p* < 0.001; log response ratio_*Ae. aegypti*_ = 0.20, CI = [0.05, 0.35], *p* = 0.01, Fig. 3). The three paired observations that evaluated mosquito burden more broadly without identifying species also found more mosquitoes in low SES neighborhoods (MEM; log response ratio = 0.92, CI = [0.40, 1.16], *p* < 0.001, Fig. 3). Overall, *Ae. albopictus*, *Ae. aegypti*, and non-distinguished mosquitoes were 62%, 22%, and 151% higher in low SES neighborhoods, respectively. *Culex* spp. were marginally more associated with low SES neighborhoods in urban cities (MEM; log response ratio = 0.64, CI = [-0.01, 1.23], *p* = 0.05, Fig. 3), but the variation around the effect magnitude was high. The overall residual heterogeneity of effect sizes was large for the mixed-effects model (Q_E_ = 74.87, df = 17, *p* < 0.001), suggesting that important unmeasured factors contribute to the effects of socioeconomic status on mosquito burden.

**Figure 3.**
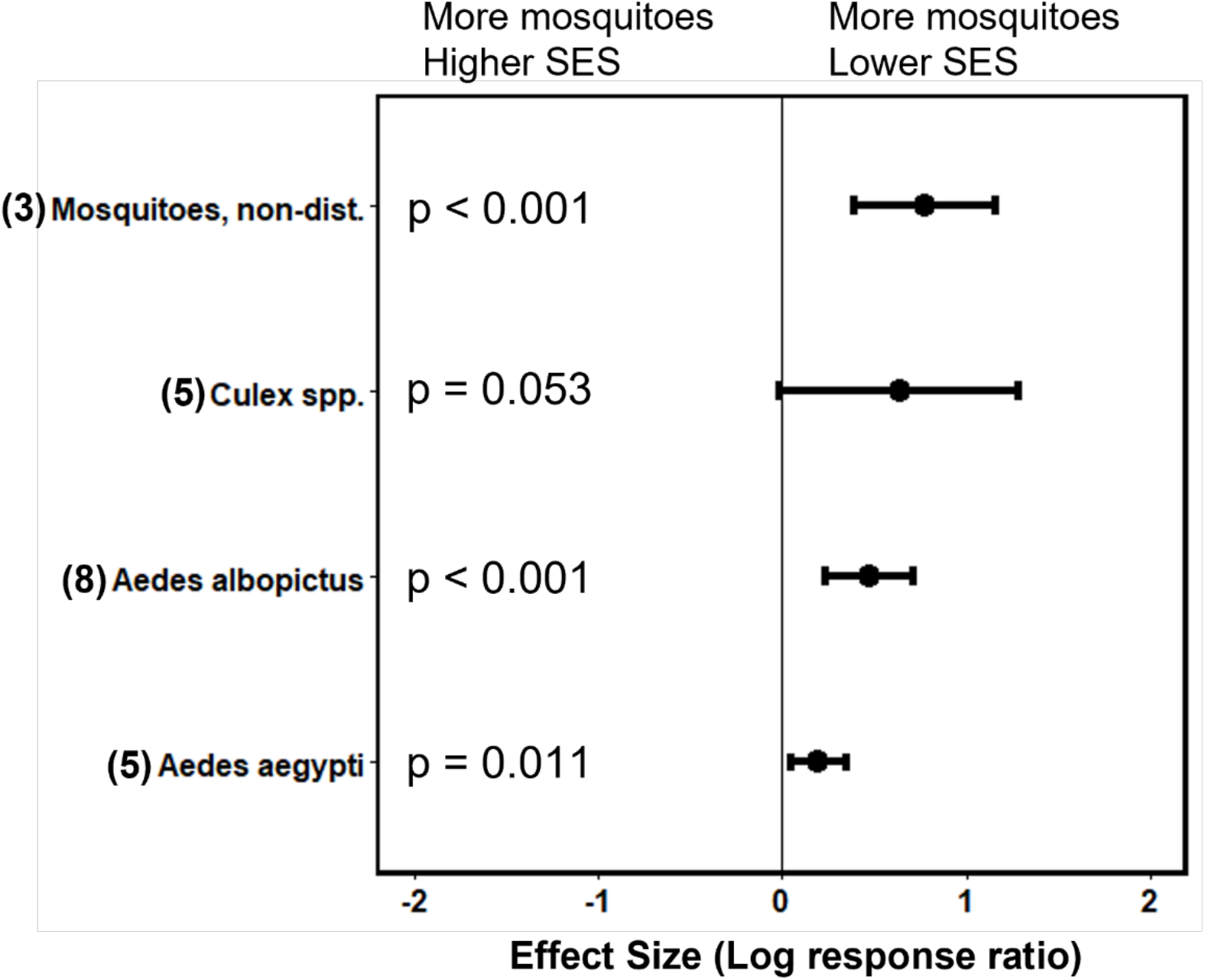
The relationship between SES and mosquito burden using mean effect sizes (log response ratio [LRR]). Positive effect sizes indicate mosquito burden was greater in low SES neighborhoods, whereas negative effect sizes denote higher mosquito burden in high SES neighborhoods. Means of LRR are shown alongside 95% CI. The number provided in the parentheses are the number of paired observations analyzed for mosquito taxon. The results are from mixed-effects models; mean effect sizes are statistically different if their 95% confidence intervals (CI) do not overlap zero.

### Study 2: Spatiotemporal patterns and spread of urban mosquitoes in Baltimore

We were able to successful predict the spatial distribution of *Ae. albopictus* populations across residential neighborhoods in Baltimore. The mean speed across all localities in Baltimore was 1.27 km year^-1^, with a median of 1.05 km year^-1^. We also related the rate of mosquito spread to latitude, longitude, median household income, normalized difference vegetation index, urban trees, and abandoned buildings. We report the estimated parameters of the spatial regression model in Table 2. On average, mosquito spread was faster across three localities in Baltimore, which depended on the available socio-ecological characteristics. We found that the median household income, tree density, vegetation index, garbage, and abandoned buildings were all significantly associated with the spread of mosquitoes.

**Table 2.** Results of spatial regression model of speeds of spread (km year^-1^) for the invasive mosquito *Aedes albopictus* in Baltimore, Maryland, including posterior means and 95 % credible intervals.

| Covariate | Beta ($\beta$) | CI low | CI high |
| --- | --- | --- | --- |
| Median household income | 0.125 | -0.230 | 0.480 |
| Tree cover | -0.017 | -0.366 | 0.320 |
| NDVI | 0.032 | -0.251 | 0.335 |
| Garbage | 0.006 | -0.211 | 0.240 |
| Abandon buildings | 0.0002 | -0.006 | 0.007 |

## DISCUSSION

Because urban environments are highly heterogenous, spatiotemporal patterns of mosquitoes and associated vector-borne diseases (e.g., West Nile, dengue, and Zika viruses) are likely affected by both social and ecological factors, with important implications for public health (Walker et al. 2018, Hernandez et al. 2019). We reveal through a meta-analysis of 15 studies from the US that there is a consistent link between lower SES and higher mosquito burden in urban environments. In particular, the mosquitoes *Ae. albopictus* and *Ae. aegypti* were strongly associated with low SES neighborhoods compared to high SES neighborhoods. Furthermore, in a spatial regression analysis of *Ae. albopictus* to model its distribution in Baltimore, we found that socio-ecological covariates supported the findings of the meta-analysis, showing that traits typically associated with low SES neighborhoods (i.e., low median household incomes, garbage, and abandoned buildings) were positively correlated with *Ae. albopictus* spread. To our knowledge, this is the first study to pair a quantitative synthesis and spatial regression analysis to understand how spatiotemporal patterns of mosquitoes correlate with SES and environmental traits in urban landscapes.

Although the meta-analysis is a coarse method to evaluate broad patterns across studies relating to urban mosquitoes and SES, the spatial regression analysis of Baltimore allowed for a more fine-grain investigation of the socio-ecological traits explaining the findings from the meta-analysis. The meta-analysis found that mosquito burden is much higher in low compared to higher SES neighborhoods across a handful of US metropolitan regions, specifically for the *Ae. albopictus* and *Ae. aegypti* mosquitoes. Based on the analysis from Baltimore, the distribution of *Ae. albopictus* through the city over 3 years correlated positively with garbage, NDVI, and abandoned buildings, whereas tree cover and median household income were negatively associated with *Ae. albopictus* spread. Thus, low SES neighborhoods in urban environments appear to have co-occurring processes that encourage mosquito populations—such as increased breeding habitats and food—which is especially apparent for *Ae. albopictus* and *Ae. aegypti*.

As anthropophilic biting insects that can breed in a variety of artificial habitats, *Ae. albopictus* and *Ae. aegypti* can thrive in places with unmanaged vegetation, garbage, dilapidated buildings, and inadequate sewage, all of which are linked to low SES neighborhoods. Artificial water-holding containers (e.g., old tires, buckets, disposable containers, etc.) can serve as breeding habitats for both *Ae. albopictus* and *Ae. aegypti*, sustaining abundant populations despite the predominance of concrete structures in urban environments. Previous research shows that increased presence of artificial containers in low SES neighborhoods can often contain more abundant, diverse mosquito communities and larvae overall (LaDeau et al. 2013, Becker et al. 2014). Moreover, unmanaged vegetation can be a positive predictor of *Ae. albopictus* abundance when coupled with high levels of building abandonment (Little et al. 2017). The availability of vegetation in the “concrete jungle” can protect mosquito larva development when summer temperatures are high, thereby increasing survivability in an otherwise harsh environment (Yang et al. 2019, Yee 2008). Unkempt vegetation can also provide food resources (e.g., plant sap, nectar, or honeydew) and resting structures for male and female mosquitoes (Zhao et al. 2020).

The significance of NDVI in our spatial model reveals that the presence of any vegetation is crucial to predicting the spread of *Ae. albopictus*, while the abundance of tree cover appears less pertinent. High SES neighborhoods generally have more tree cover in urban environments, but the vegetation is often highly managed and might adversely affect mosquito populations.

There are other human-biting mosquitoes (e.g., *Culex* spp.) that occur in urban landscapes, although the findings from the meta-analysis suggest they do not differentiate by household income as strongly as *Ae. albopictus* and *Ae. aegypti*. Some studies have also implicated Culex spp. to be urban mosquitoes with disproportionate effects in low SES neighborhoods, but the findings across the literature are inconclusive and appear context-dependent. For example, several studies have found Culex mosquitoes to be more associated with low SES neighborhoods (Chaves et al. 2011, Leisnham et al. 2014), whereas other studies have found no difference and even higher abundances in high SES neighborhoods (Dowling et al. 2013, Goodman et al. 2018). One possible reason for the variation in results is that unlike *Ae. albopictus* and *Ae. aegypti*, other Culex mosquitoes are less likely to take advantage of the abandoned buildings in impoverished neighborhoods but instead occur closer to (or in) people’s homes. Another possible explanation is that the polytypic Culex mosquito complex harbors several different species, each with a different affinity for urban environments. Such differences between species can explain our results, given that we did not separate the various mosquito species within the Culex genus.

Because the meta-analysis and spatial regression analysis are only correlational, we cannot definitively pin down the underlying mechanisms explaining the relationship between mosquito burden and SES. There are other potential explanations for why income disparity can influence mosquito burden that we could not directly assess in this research. First, mosquito control efforts and source reduction are generally concentrated in wealthier neighborhoods, potentially explaining why we found less burden in high SES neighborhoods (Tedesco et al. 2010, Biehler et al. 2018, Toju and Baba 2018). Second, local resident behavior and knowledge in low vs. high SES neighborhoods might underscore the patterns observed in this study. Previous research indicates a relationship between education level and anti-mosquito practices, such as removing water-holding containers from the yard, ultimately reducing mosquito infestations (Rochlin et al. 2011, Dowling et al. 2013a, Bodner et al. 2016). Because higher SES neighborhoods generally have high levels of education (Walker et al. 2018), and therefore likely to perform more anti-mosquito practices, differences in the behaviors of residents in high vs. low SES neighborhoods could potentially explain the results from this study.

## Conclusions

One of the unintended consequences of urban wealth disparities is that it creates environments that allow mosquito vector populations to proliferate among the economically disadvantaged (Harrigan et al. 2010). Our results highlight that mosquito burden is a complex issue in urban environments that must be understood through the dual prism of sociological and ecological factors, not just the latter. We reveal that mosquito burden is concentrated among US cities’ low SES neighborhoods, suggesting regions most vulnerable to human disease due to inadequate resources and infrastructure are also the regions that experience higher exposure to mosquitoes and associated diseases. The consequences of income-associated mosquito burden are far-reaching in urban environments (LaDeau et al. 2013, Little et al. 2017), considering that low SES neighborhoods in the US are disproportionately minority populations (i.e., Black and Hispanics). As urbanization intensifies, the subsequent increased intensity and frequency of mosquito-borne diseases (Lockaby et al. 2016) will disproportionately affect minorities. Therefore, to mitigate mosquito impacts for the localities most vulnerable to disease and ensure environmental justice, equity, and inclusion, management efforts targeting mosquito populations in low SES urban neighborhoods that incorporate fine-scale socio-ecological spatiotemporal data are required.

## ACKNOWLEDGMENTS

We thank *Ecological Applications* for supporting an Invited Feature to highlight Black scholars in applied ecology. We give special thanks to Juan Corley, Gillian Bowser, and Zsolt Silberer for their guidance and efforts in organizing this Invited Feature. We also thank Jaewoo Park for statistical advice, and Shannon LaDeau for helpful feedback on the manuscript. Lastly, we give thanks to all the authors of the original studies included in the meta-analysis, as well as Dina Fonseca, Pallavi A. Kache, and Rebeca De Jesús Crespo for their literature review suggestions. The data supporting the Baltimore study was funded by a National Science Foundation – Coupled Natural Human Systems award to Shannon LaDeau. (DEB 1211797) and the Baltimore Ecosystem Study (National Science Foundation – Long Term Ecological Research (DEB 1027188).

