## Appendix 1 for "Spatiotemporal patterns of urban mosquitoes are modulated by socioeconomic status and environmental traits in the United States"

*Ecological Applications*

Senay Yitbarek^1*^, Kelvin Chen^2^, Modeline Celestin^3^, and Matthew A. McCary^3^

^1^University of North Carolina at Chapel Hill, Department of Biology, Chapel Hill, NC 27599

^2^Rutgers University, Department of Ecology and Evolution, New Brunswick, NJ 08901

^3^Rice University, Department of BioSciences, Houston, TX 77005

**Table S1.** The screening of articles from the first round using title names and abstracts. WOS = Web of Science.

| **Study ID** | **Authors** | **Year** | **Title** | **Journal** | **Source** |
| --- | --- | --- | --- | --- | --- |
| 1 | Pooseesod et al. | 2021 | Ownership and utilization of bed nets and reasons for use or non-use of bed nets among community members at risk of malaria along the Thai-Myanmar border | Malaria Journal | WOS |
| 2 | Joyce et al. | 2021 | Forest Coverage and Socioeconomic Factors Associated with Dengue in El Salvador, 2011-2013 | Vector-borne and Zoonotic diseases | WOS |
| 3 | Jiao et al. | 2021 | How public reaction to disease information across scales and the impacts of vector control methods influence disease prevalence and control efficacy | Plos Computational Biology | WOS |
| 4 | Lubinda et al. | 2021 | Climate change and the dynamics of age-related malaria incidence in Southern Africa | Environmental Research | WOS |
| 5 | Rahman et al. | 2021 | Ecological, Social, and Other Environmental Determinants of Dengue Vector Abundance in Urban and Rural Areas of Northeastern Thailand | International Journal of Environmental Research and Public Health | WOS |
| 6 | Tassembedo et al. | 2021 | Factors associated with the use of insecticide-treated nets: analysis of the 2018 Burkina Faso Malaria Indicator Survey | Malaria Journal | WOS |
| 7 | Oliveira et al. | 2021 | Vector role and human biting activity of Anophelinae mosquitoes in different landscapes in the Brazilian Amazon | Parasites and Vectors | WOS |
| 8 | Olson et al. | 2021 | Global patterns of aegyptism without arbovirus | Plos Neglected Tropical Diseases | WOS |
| 9 | Nunez-Avellaneda et al. | 2021 | Co-Circulation of All Four Dengue Viruses and Zika Virus in Guerrero, Mexico, 2019 | Vector-borne and Zoonotic diseases | WOS |
| 10 | Rijal et al. | 2021 | Epidemiology of dengue virus infections in Nepal, 2006-2009 | Infectious Diseases of Poverty | WOS |
| 11 | Zhou et al. | 2021 | Multi-Indicator and Multistep Assessment of Malaria Transmission Risks in Western Kenya | American Journal of Tropical Medicine and Hygiene | WOS |
| 12 | Pinchoff et al. | 2021 | Use of effective lids reduces presence of mosquito larvae in household water storage containers in urban and peri-urban Zika risk areas of Guatemala, Honduras, and El Salvador | Parasites and Vectors | WOS |
| 13 | Bogale et al. | 2021 | Transcriptional heterogeneity and tightly regulated changes in gene expression during Plasmodium berghei sporozoite development | Proceedings of the National Academy of Sciences | WOS |
| 14 | Iliyasu et al. | 2021 | "A child with sickle cell disease can't live with just anyone." A mixed methods study of socio-behavioral influences and severity of sickle cell disease in northern Nigeria | Health Science Reports | WOS |
| 15 | Coutinho et al. | 2021 | Zika virus public health crisis and the perpetuation of gender inequality in Brazil | Reproductive Health | WOS |
| 16 | Mahmud et al. | 2021 | Megacities as drivers of national outbreaks: The 2017 chikungunya outbreak in Dhaka, Bangladesh | Plos Neglected Tropical Diseases | WOS |
| 17 | Larsen et al. | 2021 | Implications of Insecticide-Treated Mosquito Net Fishing in Lower Income Countries | Environmental Health Perspectives | WOS |
| 18 | Uelmen et al. | 2021 | Effects of Scale on Modeling West Nile Virus Disease Risk | American Journal of Tropical Medicine and Hygiene | WOS |
| 19 | Kirstein et al. | 2021 | Natural arbovirus infection rate and detectability of indoor female Aedes aegypti from Merida, Yucatan, Mexico | Plos Neglected Tropical Diseases | WOS |
| 20 | Athni et al. | 2021 | The influence of vector-borne disease on human history: socio-ecological mechanisms | Ecology Letters | Supplemental |
| 21 | Sallam et al. | 2021 | Systematic Review: Land Cover, Meteorological, and Socioeconomic Determinants of Aedes Mosquito Habitat for Risk Mapping | Internation Journal of Environmental Research and Public Health | Supplemental |
| 22 | Zhou et al. | 2020 | Long-lasting microbial larvicides for controlling insecticide resistant and outdoor transmitting vectors: a cost-effective supplement for malaria interventions | Infectious Diseases of Poverty | WOS |
| 23 | Lorenz et al. | 2020 | Predicting Aedes aegypti infestation using landscape and thermal features | Scientific Reports | WOS |
| 24 | Doum et al. | 2020 | Dengue Seroprevalence and Seroconversion in Urban and Rural Populations in Northeastern Thailand and Southern Laos | International Journal of Environmental Research and Public Health | WOS |
| 25 | Elaagip et al. | 2020 | Seroprevalence and associated risk factors of Dengue fever in Kassala state, eastern Sudan | Plos Neglected Tropical Diseases | WOS |
| 26 | Peterson et al. | 2020 | Amplification of pathogenic Leptospira infection with greater abundance and co-occurrence of rodent hosts across a counter-urbanizing landscape | Molecular Ecology | WOS |
| 27 | Bron et al. | 2020 | Context matters: Contrasting behavioral and residential risk factors for Lyme disease between high-incidence states in the Northeastern and Midwestern United States | Ticks and Tick-Borne Diseases | WOS |
| 28 | Yang et al. | 2020 | A deltamethrin crystal polymorph for more effective malaria control | Proceedings of the National Academy of Sciences | WOS |
| 29 | Tamari et al. | 2020 | Protective effects of Olyset (R) Net on Plasmodium falciparum infection after three years of distribution in western Kenya | Malaria Journal | WOS |
| 30 | Ngugi et al. | 2020 | Risk factors for Aedes aegypti household pupal persistence in longitudinal entomological household surveys in urban and rural Kenya | Parasites and Vectors | WOS |
| 31 | Arisco et al. | 2020 | Variation inAnophelesdistribution and predictors of malaria infection risk across regions of Madagascar | Malaria Journal | WOS |
| 32 | Ameyaw et al. | 2020 | Individual, community and region level predictors of insecticide-treated net use among women in Uganda: a multilevel analysis | Malaria Journal | WOS |
| 33 | Echaubard et al. | 2020 | Fostering social innovation and building adaptive capacity for dengue control in Cambodia: a case study | Infectious Diseases of Poverty | WOS |
| 34 | Zhao et al. | 2020 | Machine learning and dengue forecasting: Comparing random forests and artificial neural networks for predicting dengue burden at national and sub-national scales in Colombia | Plos Neglected Tropical Diseases | WOS |
| 35 | Fletcher et al. | 2020 | The Relative Role of Climate Variation and Control Interventions on Malaria Elimination Efforts in El Oro, Ecuador: A Modeling Study | Frontiers in Environmental Science | WOS |
| 36 | Sidiki et al. | 2020 | Effect of Impregnated Mosquito Bed Nets on the Prevalence of Malaria among Pregnant Women in Foumban Subdivision, West Region of Cameroon | Journal of Parasitology Research | WOS |
| 37 | Ahadiji-Dabla | 2020 | Potential Roles of Environmental and Socio-Economic Factors in the Distribution of Insecticide Resistance in Anopheles gambiae sensu lato (Culicidae: Diptera) Across Togo, West Africa | Journal of Medical Entomology | WOS |
| 38 | Dudouet et al. | 2020 | Chikungunya resurgence in the Maldives and risk for importation via tourists to Europe in 2019-2020: A GeoSentinel case series | Travel Medicine and Infectious Disease | WOS |
| 39 | Chen et al. | 2020 | Effects of natural and socioeconomic factors on dengue transmission in two cities of China from 2006 to 2017 | Science of the Total Environment | WOS |
| 40 | Madewell et al. | 2020 | Inverse association between dengue, chikungunya, and Zika virus infection and indicators of household air pollution in Santa Rosa, Guatemala: A case-control study, 2011-2018 | Plos One | WOS |
| 41 | Bonnet et al. | 2020 | Impact of a community-based intervention on Aedes aegypti and its spatial distribution in Ouagadougou, Burkina Faso | Infectious Diseases of Poverty | WOS |
| 42 | Lorenz et al. | 2020 | Remote sensing for risk mapping of Aedes aegypti infestations: Is this a practical task? | Acta Tropica | WOS |
| 43 | Rodriguez et al. | 2020 | Understanding Zika virus as an STI: findings from a qualitative study of pregnant women in the Bronx | Sexually Transmitted Infections | WOS |
| 44 | Peterson et al. | 2020 | Rodent assemblage structure reflects socioecological mosaics of counter-urbanization across post-Hurricane Katrina New Orleans | Landscape and Urban Planning | WOS |
| 45 | Berry et al. | 2020 | The origins of dengue and chikungunya viruses in Ecuador following increased migration from Venezuela and Colombia | BMC Evolutionary Biology | WOS |
| 46 | Hawaria et al. | 2020 | Effects of environmental modification on the diversity and positivity of anopheline mosquito aquatic habitats at Arjo-Dedessa irrigation development site, Southwest Ethiopia | Infectious Diseases of Poverty | WOS |
| 47 | Rudasingwa and Cho | 2020 | Determinants of the persistence of malaria in Rwanda | Malaria Journal | WOS |
| 48 | Elson et al. | 2020 | Cross-sectional study of dengue-related knowledge, attitudes and practices in Villa El Salvador, Lima, Peru | BMJ Open | WOS |
| 49 | Dinkel et al. | 2020 | Relationship of sanitation, water boiling, and mosquito nets to health biomarkers in a rural subsistence population | American Journal of Human Biology | WOS |
| 50 | Rose et al. | 2020 | Climate and Urbanization Drive Mosquito Preference for Humans | Current Biology | Supplemental |
| 51 | Whitney et al. | 2020 | Systematic review: the impact of socioeconomic factors on Aedes aegypti mosquito distribution in the mainland United States | Review Environmental Health | Supplemental |
| 52 | Ferreira et al. | 2020 | The Asian tiger mosquito in Brazil: Observations on biology and ecological interactions since its first detection in 1986 | Acta Tropica | Supplemental |
| 53 | Madewell et al. | 2019 | Associations between household environmental factors and immature mosquito abundance in Quetzaltenango, Guatemala | BMC Public Health | WOS |
| 54 | Zheng et al. | 2019 | Seasonality modeling of the distribution of Aedes albopictus in China based on climatic and environmental suitability | Infectious Diseases of Poverty | WOS |
| 55 | Adams et al. | 2019 | Water insecurity and urban poverty in the Global South: Implications for health and human biology | American Journal of Human Biology | WOS |
| 56 | Tan et al. | 2019 | A popular Indian clove-based mosquito repellent is less effective against Culex quinquefasciatus and Aedes aegypti than DEET | Plos One | WOS |
| 57 | Biehler et al. | 2019 | Knowing nature and community through mosquitoes: reframing pest management through lay vector ecologies | Local Environment | WOS |
| 58 | Malede et al. | 2019 | Barriers of persistent long-lasting insecticidal nets utilization in villages around Lake Tana, Northwest Ethiopia: a qualitative study | BMC Public Health | WOS |
| 59 | Berthe et al. | 2019 | Poverty and food security: drivers of insecticide-treated mosquito net misuse in Malawi | Malaria Journal | WOS |
| 60 | Mutua et al. | 2019 | A Qualitative Study on Gendered Barriers to Livestock Vaccine Uptake in Kenya and Uganda and Their Implications on Rift Valley Fever Control | Vaccines | WOS |
| 61 | Messina et al. | 2019 | he current and future global distribution and population at risk of dengue | Nature Microbiology | WOS |
| 62 | Ogashawara | 2019 | Spatial-Temporal Assessment of Environmental Factors Related to Dengue Outbreaks in Sao Paulo, Brazil | Geohealth | WOS |
| 63 | Iyanda | 2019 | Regional variation and demographic factors associated with knowledge of malaria risk and prevention strategies among pregnant women in Nigeria | Women and Health | WOS |
| 64 | Mouhamadou et al. | 2019 | Evidence of insecticide resistance selection in wild Anopheles coluzzii mosquitoes due to agricultural pesticide use | Infectious Diseases of Poverty | WOS |
| 65 | Parker et al. | 2019 | A Mosquito Workshop and Community Intervention: A Pilot Education Campaign to Identify Risk Factors Associated with Container Mosquitoes in San Pedro Sula, Honduras | International Journal of Environmental Research and Public Health | WOS |
| 66 | Essendi et al. | 2019 | Epidemiological risk factors for clinical malaria infection in the highlands of Western Kenya | Malaria Journal | WOS |
| 67 | Finda et al. | 2019 | Linking human behaviours and malaria vector biting risk in south-eastern Tanzania | Plos One | WOS |
| 68 | Whiteman et al. | 2019 | Aedes Mosquito Infestation in Socioeconomically Contrasting Neighborhoods of Panama City | Ecohealth | WOS |
| 69 | Kolimenaskis et al. | 2019 | On lifestyle trends, health and mosquitoes: Formulating welfare levels for control of the Asian tiger mosquito in Greece | Plos Neglected Tropical Disease | WOS |
| 70 | Beaulieu et al. | 2019 | Simplification of vector communities during suburban succession | Plos One | WOS |
| 71 | Ryan et al. | 2019 | Socio-Ecological Factors Associated with Dengue Risk and Aedes aegypti Presence in the Galapagos Islands, Ecuador | International Journal of Environmental Research and Public Health | WOS |
| 72 | Killeen et al. | 2019 | Suppression of malaria vector densities and human infection prevalence associated with scale-up of mosquito-proofed housing in Dar es Salaam, Tanzania: re-analysis of an observational series of parasitological and entomological surveys | Lancet Planetary Health | WOS |
| 73 | Moraru and Goddard | 2019 | Allergy to arthropods | Goddard Guide to Arthropods of Medical Importance | WOS |
| 74 | Koenker et al. | 2019 | Quantifying Seasonal Variation in Insecticide-Treated Net Use among Those with Access | American Journal of Tropical Medicine and Hygiene | WOS |
| 75 | Seyfarth et al. | 2019 | Five years following first detection of Anopheles stephensi (Diptera: Culicidae) in Djibouti, Horn of Africa: populations established—malaria emerging | Parasitology Research | Supplemental |
| 76 | Rael et al. | 2018 | Rat Lungworm Infection in Rodents across Post-Katrina New Orleans, Louisiana, USA | Emerging Infectious Diseases | WOS |
| 77 | Estallo et al. | 2018 | Modelling the distribution of the vector Aedes aegypti in a central Argentine city | Medical and Veterinary Entomology | WOS |
| 78 | Whiteman et al. | 2018 | Socioeconomic and demographic predictors of resident knowledge, attitude, and practice regarding arthropod-borne viruses in Panama | BMC Public Health | WOS |
| 79 | Stefopoulou et al. | 2018 | Reducing Aedes albopictus breeding sites through education: A study in urban area | Plos One | WOS |
| 80 | Juarbe-Rey et al. | 2018 | Using Risk Communication Strategies for Zika Virus Prevention and Control Driven by Community-Based Participatory Research | International Journal of Environmental Research and Public Health | WOS |
| 81 | Lee et al. | 2018 | Disparities in Zika Virus Testing and Incidence Among Women of Reproductive Age-New York City, 2016 | Journal of Public Health Management and Practice | WOS |
| 82 | Orji et al. | 2018 | Perception and Utilization of Insecticide-Treated Mosquito Net among Caregivers of Children in Abakaliki, Nigeria | Annals of African Medicine | WOS |
| 83 | Krystosik et al. | 2018 | Neighborhood Violence Impacts Disease Control and Surveillance: Case Study of Cali, Colombia from 2014 to 2016 | International Journal of Environmental Research and Public Health | WOS |
| 84 | Tabiri et al. | 2018 | The use of mesh for inguinal hernia repair in northern Ghana | Journal of Surgical Research | WOS |
| 85 | Bangert et al. | 2018 | Economic analysis of dengue prevention and case management in the Maldives | Plos Neglected Tropical Diseases | WOS |
| 86 | Stoney et al. | 2018 | Infectious diseases acquired by international travellers visiting the USA | Journal of Travel Medicine | WOS |
| 87 | Xia et al. | 2018 | Photoperiodic diapause in a subtropical population of Aedes albopictus in Guangzhou, China: optimized field-laboratory-based study and statistical models for comprehensive characterization | Infectious Diseases of Poverty | WOS |
| 88 | Khatib et al. | 2018 | Epidemiological characterization of malaria in rural southern Tanzania following China-Tanzania pilot joint malaria control baseline survey | Malaria Journal | WOS |
| 89 | Atiglo et al. | 2018 | ense of community and willingness to support malaria intervention programme in urban poor Accra, Ghana | Malaria Journal | WOS |
| 90 | Babalola et al. | 2018 | Factors associated with caregivers' consistency of use of bed nets in Nigeria: a multilevel multinomial analysis of survey data | Malaria Journal | WOS |
| 91 | Rossi et al. | 2018 | The spread of mosquito-borne viruses in modern times: A spatio-temporal analysis of dengue and chikungunya | Spatial and Spatial-Temporal Epidemiology | WOS |
| 92 | Lo lacono et al. | 2018 | Environmental limits of Rift Valley fever revealed using ecoepidemiological mechanistic models | Proceedings of the National Academy of Sciences | WOS |
| 93 | Fang et al. | 2018 | Co-circulation of Aedes flavivirus, Culex flavivirus, and Quang Binh virus in Shanghai, China | Infectious Diseases of Poverty | WOS |
| 94 | Paul et al. | 2018 | Risk factors for the presence of dengue vector mosquitoes, and determinants of their prevalence and larval site selection in Dhaka, Bangladesh | Plos One | WOS |
| 95 | Colon-Gonzalez et al. | 2018 | Limiting global-mean temperature increase to 1.5-2 degrees C could reduce the incidence and spatial spread of dengue fever in Latin America | Proceedings of the National Academy of Sciences | WOS |
| 96 | Elf et al. | 2018 | Sources of household air pollution and their association with fine particulate matter in low-income urban homes in India | Journal of Exposure Science and Environmental Epidemiology | WOS |
| 97 | Dhimal et al. | 2018 | Threats of Zika virus transmission for Asia and its Hindu-Kush Himalayan region | Infectious Diseases of Poverty | WOS |
| 98 | Robinson et al. | 2018 | Antibiotic Utilization and the Role of Suspected and Diagnosed Mosquito-borne Illness Among Adults and Children With Acute Febrile Illness in Pune, India | Clinical Infectious Diseases | WOS |
| 99 | Huber et al. | 2018 | Seasonal temperature variation influences climate suitability for dengue, chikungunya, and Zika transmission | Plos Neglected Tropical Diseases | WOS |
| 100 | Brand et al. | 2018 | A phytosociological analysis and description of wetland vegetation and ecological factors associated with locations of high mortality for the 2010-11 Rift Valley fever outbreak in South Africa | Plos One | WOS |
| 101 | Khieu et al. | 2018 | How elimination of lymphatic filariasis as a public health problem in the Kingdom of Cambodia was achieved | Infectious Diseases of Poverty | WOS |
| 102 | Amoroso et al. | 2018 | Next wave of interventions to reduce under-five mortality in Rwanda: a cross-sectional analysis of demographic and health survey data | BMC Pediatrics | WOS |
| 103 | Rek et al. | 2018 | Rapid improvements to rural Ugandan housing and their association with malaria from intense to reduced transmission: a cohort study | Lancet Planetary Health | WOS |
| 104 | Alam et al. | 2018 | Climatic changes and vulnerability of household food utilisation in Malaysian East Coast Economic Region | International Journal of Environmental and Sustainable Development | WOS |
| 105 | Howells et al. | 2018 | Zika virus in American Samoa: challenges to prevention in the context of health disparities and non-communicable disease | Annals of Human Biology | WOS |
| 106 | Piedrahita et al. | 2018 | Risk Factors Associated with Dengue Transmission and Spatial Distribution of High Seroprevalence in Schoolchildren from the Urban Area of Medellin, Colombia | Canadian Journal of Infectious Disease and Medical Microbiology | WOS |
| 107 | Das et al. | 2018 | Does involvement of local NGOs enhance public service delivery? Cautionary evidence from a malaria-prevention program in India | Health Economics | WOS |
| 108 | Heath et al. | 2018 | The Identification of Risk Factors for Chronic Chikungunya Arthralgia in Grenada, West Indies: A Cross-Sectional Cohort Study | Open Forum Infectious Diseases | WOS |
| 109 | Hendrixson et al. | 2018 | Use of a novel supplementary food and measures to control inflammation in malnourished pregnant women in Sierra Leone to improve birth outcomes: study protocol for a prospective, randomized, controlled clinical effectiveness trial | BMC Nutrition | WOS |
| 110 | Richards et al. | 2017 | Regional survey of mosquito control knowledge | Journal of the American Mosquito Control Association | WOS |
| 111 | Segal et al. | 2017 | Urinary concentrations of 3-(diethylcarbamoyl) benzoic acid (DCBA), a major metabolite of N, N-diethyl-m-toluamide (DEET) and semen parameters among men attending a fertility center | Human Reproduction | WOS |
| 112 | Chaparro et al. | 2017 | Urban malaria transmission in a non-endemic area in the Andean region of Colombia | Memorias Do Instituto Oswaldo Cruz | WOS |
| 113 | Obenauer et al. | 2017 | The importance of human population characteristics in modeling Aedes aegypti distributions and assessing risk of mosquito-borne infectious diseases | Tropical Medicine and Health | WOS |
| 114 | Ortiz et al. | 2017 | Post-earthquake Zika virus surge: Disaster and public health threat amid climatic conduciveness | Scientific Reports | WOS |
| 115 | Lana et al. | 2017 | Socioeconomic and demographic characterization of an endemic malaria region in Brazil by multiple correspondence analysis | Malaria Journal | WOS |
| 116 | Al-Jabi | 2017 | Global research trends in West Nile virus from 1943 to 2016: a bibliometric analysis | Globalization and Health | WOS |
| 117 | Krystosik et all | 2017 | Community context and sub-neighborhood scale detail to explain dengue, chikungunya and Zika patterns in Cali, Colombia | Plos One | WOS |
| 118 | Dunkel et al. | 2017 | Women's perceptions of health, quality of life, and malaria management in Kakamega County, Western Province, Kenya | Geojournal | WOS |
| 119 | Emerson and Glaser | 2017 | Cytonuclear Epistasis Controls the Density of Symbiont Wolbachia pipientis in Nongonadal Tissues of Mosquito Culex quinquefasciatus | G3-Genes Genomes Genetics | WOS |
| 120 | Alaniz et al. | 2017 | Spatial quantification of the world population potentially exposed to Zika virus | International Journal of Epidemiology | WOS |
| 121 | Marteleto et al. | 2017 | Women's Reproductive Intentions and Behaviors during the Zika Epidemic in Brazil | Population and Development Review | WOS |
| 122 | Hicks et al. | 2017 | Neurodevelopmental Delay Diagnosis Rates Are Increased in a Region with Aerial Pesticide Application | Frontiers in Pediatrics | WOS |
| 123 | Ogoma et al. | 2017 | A low technology emanator treated with the volatile pyrethroid transfluthrin confers long term protection against outdoor biting vectors of lymphatic filariasis, arboviruses and malaria | Plos Neglected Tropical Diseases | WOS |
| 124 | Fokam et al. | 2017 | Determination of the predictive factors of long-lasting insecticide-treated net ownership and utilisation in the Bamenda Health District of Cameroon | BMC Public Health | WOS |
| 125 | Reyes-Castro et al. | 2017 | Spatio-temporal and neighborhood characteristics of two dengue outbreaks in two arid cities of Mexico | Acta Tropica | WOS |
| 126 | Killeen et al. | 2017 | Developing an expanded vector control toolbox for malaria elimination | BMJ Global Health | WOS |
| 127 | Goindin et al. | 2017 | Levels of insecticide resistance to deltamethrin, malathion, and temephos, and associated mechanisms in Aedes aegypti mosquitoes from the Guadeloupe and Saint Martin islands (French West Indies) | Infectious Diseases of Poverty | WOS |
| 128 | Moreno-Madrinan and Turell | 2017 | Factors of Concern Regarding Zika and Other Aedes aegypti-Transmitted Viruses in the United States | Journal of Medical Entomology | WOS |
| 129 | Heydari et al. | 2017 | Household Dengue Prevention Interventions, Expenditures, and Barriers to Aedes aegypti Control in Machala, Ecuador | International Journal of Enviromental Research and Public Health | WOS |
| 130 | Anaya et al. | 2017 | A comprehensive analysis and immunobiology of autoimmune neurological syndromes during the Zika virus outbreak in Cucuta, Colombia | Journal of Autoimmunity | WOS |
| 131 | Weaver et al. | 2017 | Pilot Intervention Study of Household Ventilation and Fine Particulate Matter Concentrations in a Low-Income Urban Area, Dhaka, Bangladesh | American Journal of Tropical Medicine and Hygiene | WOS |
| 132 | Legorreta-Soberanis et al. | 2017 | Household costs for personal protection against mosquitoes: secondary outcomes from a randomised controlled trial of dengue prevention in Guerrero state, Mexico | BMC Public Health | WOS |
| 133 | Choo et al. | 2017 | School-based health education in Yucatan, Mexico about the Chikungunya virus and mosquito illness prevention | Infectious Disease Reports | WOS |
| 134 | Ernst et al. | 2017 | Aedes aegypti (Diptera: Culicidae) Longevity and Differential Emergence of Dengue Fever in Two Cities in Sonora, Mexico | Journal of Medical Entomology | WOS |
| 135 | Laaksonen et al. | 2017 | Filarioid nematodes, threat to arctic food safety and security | Game Meat Hygiene: Food Safety and Security | WOS |
| 136 | Singer | 2017 | The spread of Zika and the potential for global arbovirus syndemics | Global Public Health | WOS |
| 137 | Wood et al. | 2017 | Human infectious disease burdens decrease with urbanization but not with biodiversity | Philosophical Transactions of the Royal Society B: Biological Sciences | Supplemental |
| 138 | Santos-Vega | 2016 | Population Density, Climate Variables and Poverty Synergistically Structure Spatial Risk in Urban Malaria in India | Plos Neglected Tropical Diseases | WOS |
| 139 | Delmelle et al. | 2016 | A spatial model of socioeconomic and environmental determinants of dengue fever in Cali, Colombia | Acta Tropica | WOS |
| 140 | Nyamwaya et al. | 2016 | Detection of West Nile virus in wild birds in Tana River and Garissa Counties, Kenya | BMC Infectious Diseases | WOS |
| 141 | Brown et al. | 2016 | Household perceptions and subjective valuations of indoor residual spraying programmes to control malaria in northern Uganda | Infectious Diseases of Poverty | WOS |
| 142 | Aung et al. | 2016 | Ownership and Use of Insecticide-Treated Nets among People Living in Malaria Endemic Areas of Eastern Myanmar | Plos One | WOS |
| 143 | Higuera-Mendieta et al. | 2016 | KAP Surveys and Dengue Control in Colombia: Disentangling the Effect of Sociodemographic Factors Using Multiple Correspondence Analysis | Plos Neglected Tropical Diseases | WOS |
| 144 | Samy et al. | 2016 | Mapping the global geographic potential of Zika virus spread | Memorias Do Instituto Oswaldo Cruz | WOS |
| 145 | Wiljayanti et al. | 2016 | The Importance of Socio-Economic Versus Environmental Risk Factors for Reported Dengue Cases in Java, Indonesia | Plos Neglected Tropical Diseases | WOS |
| 146 | Tusting et al. | 2016 | Why is malaria associated with poverty? Findings from a cohort study in rural Uganda | Infectious Diseases of Poverty | WOS |
| 147 | Mondal et al. | 2016 | Efficacy, Safety and Cost of Insecticide Treated Wall Lining, Insecticide Treated Bed Nets and Indoor Wall Wash with Lime for Visceral Leishmaniasis Vector Control in the Indian Sub-continent: A Multi-country Cluster Randomized Controlled Trial | Plos Neglected Tropical Diseases | WOS |
| 148 | van Elijk et al. | 2016 | The use of mosquito repellents at three sites in India with declining malaria transmission: surveys in the community and clinic | Parasites and Vectors | WOS |
| 149 | Paploski et al. | 2016 | Storm drains as larval development and adult resting sites for Aedes aegypti and Aedes albopictus in Salvador, Brazil | Parasites and Vectors | WOS |
| 150 | Birkett | 2016 | Status of vaccine research and development of vaccines for malaria | Vaccines | WOS |
| 151 | Mwangungulu et al. | 2016 | Crowdsourcing Vector Surveillance: Using Community Knowledge and Experiences to Predict Densities and Distribution of Outdoor-Biting Mosquitoes in Rural Tanzania | Plos One | WOS |
| 152 | Kuan et al. | 2016 | Seroprevalence of Anti-Chikungunya Virus Antibodies in Children and Adults in Managua, Nicaragua, After the First Chikungunya Epidemic, 2014-2015 | Plos Neglected Tropical Diseases | WOS |
| 153 | Gil et al. | 2016 | Spatial spread of dengue in a non-endemic tropical city in northern Argentina | Acta Tropica | WOS |
| 154 | Ngonghala et al. | 2016 | Interplay between insecticide-treated bed-nets and mosquito demography: implications for malaria control | Journal of Theoretical Biology | WOS |
| 155 | Bodner et al. | 2016 | Effectiveness of Print Education at Reducing Urban Mosquito Infestation through Improved Resident-Based Management | Plos One | WOS |
| 156 | Gyawali et al. | 2016 | The global spread of Zika virus: is public and media concern justified in regions currently unaffected? | Infectious Diseases of Poverty | WOS |
| 157 | Kaindoa et al. | 2016 | Correlations between household occupancy and malaria vector biting risk in rural Tanzanian villages: implications for high-resolution spatial targeting of control interventions | Malaria Journal | WOS |
| 158 | Moon et al. | 2016 | Factors associated with the use of mosquito bed nets: results from two cross-sectional household surveys in Zambezia Province, Mozambique | Malaria Journal | WOS |
| 159 | Dickinson et al. | 2016 | Willingness to Pay for Mosquito Control in Key West, Florida and Tucson, Arizona | American Journal of Tropical Medicine and Hygiene | WOS |
| 160 | Chitunhu and Musenge | 2016 | Spatial and socio-economic effects on malaria morbidity in children under 5 years in Malawi in 2012 | Spatial and Spatial-Temporal Epidemiology | WOS |
| 161 | Rosas-Aguirre et al. | 2016 | Epidemiology of Plasmodium vivax Malaria in Peru | American Journal of Tropical Medicine and Hygiene | WOS |
| 162 | Bizimana et al. | 2016 | Modelling homogeneous regions of social vulnerability to malaria in Rwanda | Geospatial Health | WOS |
| 163 | Nankabirwa et al. | 2015 | Estimating malaria parasite prevalence from community surveys in Uganda: a comparison of microscopy, rapid diagnostic tests and polymerase chain reaction | Malaria Journal | WOS |
| 164 | Mmbando et al. | 2015 | Effects of a new outdoor mosquito control device, the mosquito landing box, on densities and survival of the malaria vector, Anopheles arabiensis, inside controlled semi-field settings | Malaria Journal | WOS |
| 165 | Londono-Renteria et al. | 2015 | Aedes aegypti anti-salivary gland antibody concentration and dengue virus exposure history in healthy individuals living in an endemic area in Colombia | Biomedica | WOS |
| 166 | Ryan et al. | 2015 | Mapping Physiological Suitability Limits for Malaria in Africa Under Climate Change | Vector-borne and Zoonotic diseases | WOS |
| 167 | Paz-Soldan et al. | 2015 | Dengue Knowledge and Preventive Practices in Iquitos, Peru | American Journal of Tropical Medicine and Hygiene | WOS |
| 168 | Mathanga et al. | 2015 | The effectiveness of long-lasting, insecticide-treated nets in a setting of pyrethroid resistance: a case-control study among febrile children 6 to 59 months of age in Machinga District, Malawi | Malaria Journal | WOS |
| 169 | Biadgillign et al. | 2015 | Determinants of willingness to pay for the retreatment of insecticide treated mosquito nets in rural area of eastern Ethiopia | International Journal for Equity in Health | WOS |
| 170 | Salmon-Mulanovich et al. | 2015 | Economic Burden of Dengue Virus Infection at the Household Level among Residents of Puerto Maldonado, Peru | American Journal of Tropical Medicine and Hygiene | WOS |
| 171 | Mathanga et al. | 2015 | The High Burden of Malaria in Primary School Children in Southern Malawi | American Journal of Tropical Medicine and Hygiene | WOS |
| 172 | Shabani et al. | 2015 | Knowledge, attitudes and practices on Rift Valley fever among agro pastoral communities in Kongwa and Kilombero districts, Tanzania | BMC Infectious Diseases | WOS |
| 173 | Gomez-Perez et al. | 2015 | Controlled human malaria infection by intramuscular and direct venous inoculation of cryopreserved Plasmodium falciparum sporozoites in malaria-naive volunteers: effect of injection volume and dose on infectivity rates | Malaria Journal | WOS |
| 174 | LaDeau et al. | 2015 | The ecological foundations of transmission potential and vector-borne disease in urban landscapes | Functional Ecology | WOS |
| 175 | Acevedo et al. | 2015 | patial Heterogeneity, Host Movement and Mosquito-Borne Disease Transmission | Plos One | WOS |
| 176 | Liu et al. | 2015 | Coverage, use and maintenance of bed nets and related influence factors in Kachin Special Region II, northeastern Myanmar | Malaria Journal | WOS |
| 177 | Feldstein et al. | 2015 | Dengue on islands: a Bayesian approach to understanding the global ecology of dengue viruses | Transactions of the Royal Society of Tropical Medicine and Hygiene | WOS |
| 178 | Lai et al. | 2015 | The changing epidemiology of dengue in China, 1990-2014: a descriptive analysis of 25 years of nationwide surveillance data | BMC Medicine | WOS |
| 179 | Mayala et al. | 2015 | Knowledge, perception and practices about malaria, climate change, livelihoods and food security among rural communities of central Tanzania | Infectious Diseases of Poverty | WOS |
| 180 | Obaldia | 2015 | Determinants of low socio-economic status and risk of Plasmodium vivax malaria infection in Panama (2009-2012): a case-control study | Malaria Journal | WOS |
| 181 | Oyekale | 2015 | Do ownership of mosquito nets, dwelling characteristics and mothers’ socio-economic status influence malaria morbidity among children under the age of 5 in Cameroon? | International Journal of Occupational Medicine and Environmental Health | WOS |
| 182 | Chiu et al. | 2014 | A Probabilistic Spatial Dengue Fever Risk Assessment by a Threshold-Based-Quantile Regression Method | Plos One | WOS |
| 183 | Wang | 2014 | Prevention measures and socio-economic development result in a decrease in malaria in Hainan, China | Malaria Journal | WOS |
| 184 | Noden et al. | 2014 | Risk assessment of flavivirus transmission in Namibia | Acta Tropica | WOS |
| 185 | Sangoro et al. | 2014 | A cluster-randomized controlled trial to assess the effectiveness of using 15% DEET topical repellent with long-lasting insecticidal nets (LLINs) compared to a placebo lotion on malaria transmission | Malaria Journal | WOS |
| 186 | Guagliardo et al. | 2014 | Patterns of Geographic Expansion of Aedes aegypti in the Peruvian Amazon | Plos Neglected Tropical Diseases | WOS |
| 187 | Ruiz et al. | 2014 | Implementation of Malaria Dynamic Models in Municipality Level Early Warning Systems in Colombia. Part I: Description of Study Sites | American Journal of Tropical Medicine and Hygiene | WOS |
| 188 | Guerra et al. | 2014 | A global assembly of adult female mosquito mark-release-recapture data to inform the control of mosquito-borne pathogens | Parasites and Vectors | WOS |
| 189 | Wang et al. | 2014 | Factors influencing US canine heartworm (Dirofilaria immitis) prevalence | Parasites and Vectors | WOS |
| 190 | Chang et al. | 2014 | Social Justice, Climate Change, and Dengue | Health and Human Rights | WOS |
| 191 | Blake and Garcia-Blanco | 2014 | Human Genetic Variation and Yellow Fever Mortality during 19th Century US Epidemics | MBIO | WOS |
| 192 | Attaway et al. | 2014 | Assessing the methods needed for improved dengue mapping: a SWOT analysis | Pan African Medical Journal | WOS |
| 193 | Karunamoorthi and Hailu | 2014 | Insect repellent plants traditional usage practices in the Ethiopian malaria epidemic-prone setting: an ethnobotanical survey | Journal of Ethnobiology and Ethnomedicine | WOS |
| 194 | Liu et al. | 2014 | Is Housing Quality Associated with Malaria Incidence among Young Children and Mosquito Vector Numbers? Evidence from Korogwe, Tanzania | Plos One | WOS |
| 195 | Sezi et al. | 2014 | The phenomenon of diminishing -returns in the use of bed nets and indoor house spraying and the emerging place of antimalarial medicines in the control of malaria in Uganda | African Health Sciences | WOS |
| 196 | Brown et al. | 2014 | Human impacts have shaped historical and recent evolution in Aedes aegypti, the dengue and yellow fever mosquito | Evolution | Supplemental |
| 197 | Bradley et al. | 2013 | Reduced Prevalence of Malaria Infection in Children Living in Houses with Window Screening or Closed Eaves on Bioko Island, Equatorial Guinea | Plos One | WOS |
| 198 | Ibarra et al. | 2013 | Dengue Vector Dynamics (Aedes aegypti) Influenced by Climate and Social Factors in Ecuador: Implications for Targeted Control | Plos One | WOS |
| 199 | Moise et al. | 2013 | Geographic Assessment of Unattended Swimming Pools in Post-Katrina New Orleans, 2006-2008 | Annals of the Association of American Geographers | WOS |
| 200 | Hagenlocher et al. | 2013 | Assessing socioeconomic vulnerability to dengue fever in Cali, Colombia: statistical vs expert-based modeling | International Journal of Health Geographics | WOS |
| 201 | Chen-Hussey et al. | 2013 | Can Topical Insect Repellents Reduce Malaria? A Cluster-Randomised Controlled Trial of the Insect Repellent N,N-diethyl-m-toluamide (DEET) in Lao PDR | Plos One | WOS |
| 202 | Njau et al. | 2013 | Exploring the impact of targeted distribution of free bed nets on households bed net ownership, socio-economic disparities and childhood malaria infection rates: analysis of national malaria survey data from three sub-Saharan Africa countries | Malaria Journal | WOS |
| 203 | Geraghty et al. | 2013 | Correlation Between Aerial Insecticide Spraying to Interrupt West Nile Virus Transmission and Emergency Department Visits in Sacramento County, California | Public Health Reports | WOS |
| 204 | Das et al. | 2013 | Community perceptions on malaria and care-seeking practices in endemic Indian settings: policy implications for the malaria control programme | Malaria Journal | WOS |
| 205 | Okumu et al. | 2013 | Mathematical evaluation of community level impact of combining bed nets and indoor residual spraying upon malaria transmission in areas where the main vectors are Anopheles arabiensis mosquitoes | Parasites and Vectors | WOS |
| 206 | Bashar et al. | 2012 | Socio-demographic factors influencing knowledge, attitude and practice (KAP) regarding malaria in Bangladesh | BMC Public Health | WOS |
| 207 | Clasen et al. | 2012 | The effect of improved rural sanitation on diarrhoea and helminth infection: design of a cluster-randomized trial in Orissa, India | Emerging Themes in Epidemiology | WOS |
| 208 | Bennett et al. | 2012 | Household Possession and Use of Insecticide-Treated Mosquito Nets in Sierra Leone 6 Months after a National Mass-Distribution Campaign | Plos One | WOS |
| 209 | Hsueh et al. | 2012 | Spatio-temporal patterns of dengue fever cases in Kaoshiung City, Taiwan, 2003-2008 | Applied Geography | WOS |
| 210 | Booth et al. | 2012 | Molecular Markers Reveal Infestation Dynamics of the Bed Bug (Hemiptera: Cimicidae) Within Apartment Buildings | Journal of Medical Entomology | WOS |
| 211 | Khatib et al. | 2012 | Routine delivery of artemisinin-based combination treatment at fixed health facilities reduces malaria prevalence in Tanzania: an observational study | Malaria Journal | WOS |
| 212 | Welch and Fuster | 2012 | Barriers in access to insecticide-treated bednets for malaria prevention: An analysis of Cambodian DHS data | Journal of Vector Borne Diseases | WOS |
| 213 | Mwangangi et al. | 2012 | Mosquito species abundance and diversity in Malindi, Kenya and their potential implication in pathogen transmission | Parasitology Research | WOS |
| 214 | Graves et al. | 2011 | Factors associated with mosquito net use by individuals in households owning nets in Ethiopia | Malaria Journal | WOS |
| 215 | Mutuku et al. | 2011 | Impact of insecticide-treated bed nets on malaria transmission indices on the south coast of Kenya | Malaria Journal | WOS |
| 216 | Garcia-Basteiro et al. | 2011 | Determinants of bed net use in children under five and household bed net ownership on Bioko Island, Equatorial Guinea | Malaria Journal | WOS |
| 217 | Cordeiro et al. | 2011 | Spatial distribution of the risk of dengue fever in southeast Brazil, 2006-2007 | BMC Public Health | WOS |
| 218 | Delgado-Petrocelli et al. | 2011 | Geospatial tools for the identification of a malaria corridor in Estado Sucre, a Venezuelan north-eastern state | Geospatial Health | WOS |
| 219 | Ngondi et al. | 2011 | Which nets are being used: factors associated with mosquito net use in Amhara, Oromia and Southern Nations, Nationalities and Peoples' Regions of Ethiopia | Malaria Journal | WOS |
| 220 | Larsen et al. | 2011 | Comparison of Lives Saved Tool model child mortality estimates against measured data from vector control studies in sub-Saharan Africa | BMC Public Health | WOS |
| 221 | Cetin et al. | 2011 | Larvicidal activity of selected plant hydrodistillate extracts against the house mosquito, Culex pipiens, a West Nile virus vector | Parasitology Research | WOS |
| 222 | Lowe et al. | 2011 | Spatio-temporal modelling of climate-sensitive disease risk: Towards an early warning system for dengue in Brazil | Computers and Geosciences | WOS |
| 223 | Ghosh and Guha | 2011 | Using a neural network for mining interpretable relationships of West Nile risk factors | Social Science and Medicine | WOS |
| 224 | Parham et al. | 2011 | Understanding and Modelling the Impact of Climate Change on Infectious Diseases - Progress and Future Challenges | Climate Change Socioeconomic Effects | WOS |
| 225 | Erickson et al. | 2010 | A dengue model with a dynamic Aedes albopictus vector population | Ecological Modelling | WOS |
| 226 | Harrigan et al. | 2010 | Economic Conditions Predict Prevalence of West Nile Virus | Plos One | WOS |
| 227 | Gitonga et al. | 2010 | Implementing school malaria surveys in Kenya: towards a national surveillance system | Malaria Journal | WOS |
| 228 | Ghosh and Guha | 2010 | Use of genetic algorithm and neural network approaches for risk factor selection: A case study of West Nile virus dynamics in an urban environment | Computers Environment and Urban Systems | WOS |
| 229 | Komatsu et al. | 2010 | Lives saved by Global Fund-supported HIV/AIDS, tuberculosis and malaria programs: estimation approach and results between 2003 and end-2007 | BMC Infectious Diseases | WOS |
| 230 | Boulanger et al. | 2010 | A Europe-South America network for climate change assessment and impact studies | Climatic Change | WOS |
| 231 | Pullan et al. | 2010 | Plasmodium infection and its risk factors in eastern Uganda | Malaria Journal | WOS |
| 232 | Onwujekwe et al. | 2009 | Are there geographic and socio-economic differences in incidence, burden and prevention of malaria? A study in southeast Nigeria | International Journal for Equity in Health | WOS |
| 233 | Steketee and Eisele | 2009 | Is the Scale Up of Malaria Intervention Coverage Also Achieving Equity? | Plos One | WOS |
| 234 | Karunamoorthi et al. | 2009 | Ethnobotanical survey of knowledge and usage custom of traditional insect/mosquito repellent plants among the Ethiopian Oromo ethnic group | Journal of Ethnopharmacology | WOS |
| 235 | Boakye et al. | 2009 | Patterns of household insecticide use and pyrethroid resistance in Anopheles gambiae sensu stricto (Diptera: Culicidae) within the Accra metropolis of Ghana | African Entomology | WOS |
| 236 | Hanson et al. | 2009 | Household ownership and use of insecticide treated nets among target groups after implementation of a national voucher programme in the United Republic of Tanzania: plausibility study using three annual cross sectional household surveys | British Medical Journal | WOS |
| 237 | Karunamoorthi et al. | 2009 | Assessment of knowledge and usage custom of traditional insect/mosquito repellent plants in Addis Zemen Town, South Gonder, North Western Ethiopia | Journal of Ethnopharmacology | WOS |
| 238 | Richie and Parekh | 2009 | Malaria | Vaccines for Biodefense and Emerging and Neglected Diseases | WOS |
| 239 | Kumar and Ramaiah | 2008 | Usage of personal-protection measures against mosquitoes and the low prevalences of Wuchereria bancrofti microfilaraemia in the Indian city of Chennai | Annals of Tropical Medcine and Parasitology | WOS |
| 240 | Swaddle and Calos | 2008 | Increased Avian Diversity Is Associated with Lower Incidence of Human West Nile Infection: Observation of the Dilution Effect | Plos One | WOS |
| 241 | Thwing et al. | 2008 | Insecticide-treated net ownership and usage in Niger after a nationwide integrated campaign | Tropical Medicine and International Health | WOS |
| 242 | Kolaczinski et al. | 2008 | Risk factors of visceral leishmaniasis in East Africa: a case-control study in Pokot territory of Kenya and Uganda | International Journal of Epidemiology | WOS |
| 243 | Somi et al. | 2007 | Is there evidence for dual causation between malaria and socioeconomic status? Findings from rural Tanzania | American Journal of Tropical Medicine and Hygiene | WOS |
| 244 | Banda et al. | 2007 | Water handling, sanitation and defecation practices in rural southern India: a knowledge, attitudes and practices study | Transactions of the Royal Society of Tropical Medicine and Hygiene | WOS |
| 245 | Howard et al. | 2007 | Malaria mosquito control using edible fish in western Kenya: preliminary findings of a controlled study | BMC Public Health | WOS |
| 246 | Agha et al. | 2007 | The impact of a hybrid social marketing intervention on inequities in access, ownership and use of insecticide-treated nets | Malaria Journal | WOS |
| 247 | Sanjana et al. | 2006 | Survey of community knowledge, attitudes, and practices during a malaria epidemic in central Java, Indonesia | American Journal of Tropical Medicine and Hygiene | WOS |
| 248 | Mathanga et al. | 2006 | Socially marketed insecticide-treated nets effectively reduce Plasmodium infection and anaemia among children in urban Malawi | Tropical Medicine and International Health | WOS |
| 249 | Keating et al. | 2005 | Self-reported malaria and mosquito avoidance in relation to household risk factors in a Kenyan coastal city | Journal of Biosocial Science | WOS |
| 250 | Grabowsky et al. | 2005 | Integrating insecticide-treated bednets into a measles vaccination campaign achieves high, rapid and equitable coverage with direct and voucher-based methods | Tropical Medicine and International Health | WOS |
| 251 | Mwenesi et al. | 2005 | Social science research in malaria prevention, management and control in the last two decades: An overview | Acta Tropica | WOS |
| 252 | Grabowsky et al. | 2005 | Distributing insecticide-treated bednets during measles vaccination: a low-cost means of achieving high and equitable coverage | Bulletin of the World Health Organization | WOS |
| 253 | Raimondo | 2005 | 'AIDS capital of the world': Representing race, sex and space in Belle Glade, Florida | Gender Place and Culture | WOS |
| 254 | Kumar et al. | 2005 | Management of filariasis using prediction rules derived from data mining | Bioinformation | WOS |
| 255 | Su and Mulla | 2004 | Documentation of high-level Bacillus sphaericus 2362 resistance in field populations of Culex quinquefasciatus breeding in polluted water in Thailand | Journal of the American Mosquito Control Association | WOS |
| 256 | Carlson et al. | 2004 | Field assessments in western Kenya link malaria vectors to environmentally disturbed habitats during the dry season | BMC Public Health | WOS |
| 257 | Alaii et al. | 2003 | Factors affecting use of permethrin-treated bed nets during a randomized controlled trial in western Kenya | American Journal of Tropical Medicine and Hygiene | WOS |
| 258 | Keating et al. | 2003 | A geographic sampling strategy for studying relationships between human activity and malaria vectors in urban Africa | American Journal of Tropical Medicine and Hygiene | WOS |
| 259 | Mulla et al. | 2003 | Emergence of resistance and resistance management in field populations of tropical Culex quinquefasciatus to the microbial control agent Bacillus sphaericus | Journal of the American Mosquito Control Association | WOS |
| 260 | Mulla et al. | 2001 | Mosquito larval control with Bacillus sphaericus: reduction in adult populations in low-income communities in Nonthaburi Province, Thailand | Journal of Vector Ecology | WOS |
| 261 | Mulla et al. | 2001 | Mosquito burden and impact on the poor: Measures and costs for personal protection in some communities in Thailand | Journal of the American Mosquito Control Association | WOS |
| 262 | Kroeger et al. | 1997 | The contribution of repellent soap to malaria control | American Journal of Tropical Medicine and Hygiene | WOS |
| 263 | Cunningham | 1997 | Lymphatic filariasis in immigrants from developing countries | American Family Physician | WOS |
| 264 | Torres | 1997 | Impact of an outbreak of dengue fever: A case study from rural Puerto Rico | Human Organization | WOS |
| 265 | Thompson et al. | 1996 | Geographical perspectives on bednet use and malaria transmission in the Gambia, west Africa | Social Science and Medicine | WOS |
| 266 | Grey | 1992 | Syphilis and AIDS in Belle-Glade, Florida 1942 and 1992 | Annals of Internal Medicine | WOS |

| **Study ID** | **Authors** | **Year** | **Title** | **Journal** | **Volume(issue): Pages** | **Reason for exclusion** | **Source** |
| --- | --- | --- | --- | --- | --- | --- | --- |
| 122 | Vorobeichik and Bergman | 2020 | Bait-Lamina Test in the Assessment of Polluted Soils: Choice of Exposure Duration | Russian Journal of Ecology | 51(5): 430-439 | No exclusion of arthropods/invertebrates; decomposition/C/N not measured | WOS |
| 123 | Toth and Hornung | 2020 | Taxonomic and Functional Response of Millipedes (Diplopoda) to Urban Soil Disturbance in a Metropolitan Area | Insects | 11(1): 25 | No exclusion of arthropods/invertebrates; decomposition/C/N not measured | WOS |
| 124 | Borgstrom et al. | 2020 | Below-ground herbivory mitigates biomass loss from above-ground herbivory of nitrogen fertilized plants | Scientific Reports | 10(1): 12752 | No exclusion of arthropods/invertebrates; decomposition/C/N not measured | WOS |
| 125 | Kristensen et al. | 2020 | Below-ground responses to insect herbivory in ecosystems with woody plant canopies: A meta-analysis | Journal of Ecology | 108(3): 917-930 | Review/meta-analysis | WOS |
| 126 | Asselman | 2020 | Does metal pollution affect the stoichiometry of soil-litter food webs? | Pedobiologia | 80: 150649 | No invertebrate exclusion or evaluation of Decomposition/C/N | WOS |
| 127 | Dourado et al. | 2020 | Ecological indices of phytophagous Hemiptera and their natural enemies on Acacia auriculiformis (Fabales: Fabaceae) plants with or without dehydrated sewage sludge application in a degraded area | PLoS ONE | 15(8): e0237261 | No exclusion of arthropods/invertebrates; decomposition/C/N not measured | WOS |
| 128 | Duval et al. | 2020 | Effects of the Gold King Mine Spill on Metal Cycling through River and Riparian Biota | Wetlands | 40(5): 1033–1046 | Aquatic system | WOS |
| 129 | Wilfahrt et al. | 2020 | Initial richness, consumer pressure and soil resources jointly affect plant diversity and resource strategies during a successional field experiment | Journal of Ecology | 108(6): 2352-2365 | No invertebrate exclusion or evaluation of Decomposition/C/N | WOS |
| 130 | Thakur et al. | 2020 | Invasive earthworms reduce chemical defense and increase herbivory and pathogen infection in native trees | Journal of Ecology | In press (DOI 10.1111/1365-2745.13504) | Decomposition/C/N not measured | WOS |
| 131 | Peng et al. | 2020 | Landscape configuration and habitat complexity shape arthropod assemblage in urban parks | Scientific Reports | 10(1): 16043 | No exclusion of arthropods/invertebrates; decomposition/C/N not measured | WOS |
| 132 | Fujii et al. | 2020 | Living litter: Dynamic trait spectra predict fauna composition | Trends in Ecology & Evolution | 35(10): 886-896 | Review/meta-analysis | WOS |
| 133 | Andrews and Ruess | 2020 | Microarthropod abundance and community structure along a chronosequence within the Tanana River floodplain, Alaska | Ecoscience | 27(4): 235-253 | No exclusion of arthropods/invertebrates; decomposition/C/N not measured | WOS |
| 134 | Prather et al. | 2020 | Micronutrients enhance macronutrient effects in a meta-analysis of grassland arthropod abundance | Global Ecology and Biogeography | 29(12): 2273-2288 | Review/meta-analysis | WOS |
| 135 | Agathokleous et al. | 2020 | Ozone affects plant, insect, and soil microbial communities: A threat to terrestrial ecosystems and biodiversity | Science Advances | 6(33): EABC1176 | No exclusion of arthropods/invertebrates; decomposition/C/N not measured | WOS |
| 136 | Liu et al. | 2020 | Relationships between plant diversity and soil microbial diversity vary across taxonomic groups and spatial scales | Ecosphere | 11(1): e02999 | Review/meta-analysis | WOS |
| 137 | Bach et al. | 2020 | Soil Biodiversity Integrates Solutions for a Sustainable Future | Sustainability | 12(7): 2662 | Review/meta-analysis | WOS |
| 138 | Heydari et al. | 2020 | Spatio-temporal heterogeneity differently drives the diversity of various trophic guilds of mesofauna in semi-arid oak forests | Trees | In press (DOI 10.1007/s00468-020-02025-3) | No exclusion of arthropods/invertebrates; decomposition/C/N not measured | WOS |
| 139 | Rocha et al. | 2020 | Stochastic and deterministic processes differently affect the community structure of edaphic mites (Acari: Mesostigmata) in the southern Brazilian Atlantic Forest | Systematic and Applied Acarology | 25(3): 577-592 | No Decomposition/C/N measured and no invertebrate exclusion | WOS |
| 140 | Siedl et al. | 2020 | Temporary non-crop habitats within arable fields: The effects of field defects on carabid beetle assemblages | Agriculture, Ecosystems & Environment | 293: 106856 | No invertebrate exclusion or evaluation of Decomposition/C/N | WOS |
| 141 | Brand et al. | 2020 | The influence of fire and other environmental factors on terrestrial gastropod species composition in an oak-hickory woodland of west-central Illinois | American Malacological Bulletin | 38(1): 39-49 | No exclusion of arthropods/invertebrates; decomposition/C/N not measured | WOS |
| 142 | Monteiro et al. | 2020 | The mistletoe Struthanthus flexicaulis reduces dominance and increases diversity of plants in campo rupestre | Flora | 271: 151690 | No exclusion of arthropods/invertebrates; decomposition/C/N not measured | WOS |
| 143 | Raymond-Leonard | 2019 | A novel set of traits to describe Collembola mouthparts: taking a bite out of the broad chewing mandible classification | Soil Biology and Biochemistry | 138: 107608 | Decomposition/C/N not measured | WOS |
| 144 | Schappe et al. | 2019 | Co-occurring fungal functional groups respond differently to tree neighborhoods and soil properties across three tropical rainforests in Panama | Microbial Ecology | 79(3): 675-685 | Decomposition/C/N not measured; effects of invertebrates not studied | WOS |
| 145 | Harris et al. | 2019 | Decline in beetle abundance and diversity in an intact temperate forest linked to climate warming | Biological Conservation | 240: 108219 | Decomposition/C/N not measured | WOS |
| 146 | Barratt et al. | 2019 | The effect of fire on terrestrial amphipods (Crustacea: Amphipoda) in a natural grassland community | Pedobiologia | 77: 150590 | Decomposition/C/N not measured | WOS |
| 147 | Stam et al. | 2019 | Cross-seasonal legacy effects of arthropod community on plant fitness in perennial plants | Journal of Ecology | 107(5): 2451-2463 | Decomposition/C/N not measured | WOS |
| 148 | Bernaola and Stout | 2019 | Effects of arbuscular mycorrhizal fungi on rice-herbivore interactions are soil-dependent | Scientific Reports | 9(1): 14037 | No exclusion of arthropods/invertebrates; decomposition/C/N not measured | WOS |
| 149 | Duran et al. | 2019 | Wildfires decrease the local-scale ecosystem spatial variability of Pinus canariensis forests during the first two decades post fire | International Journal of Wildland Fire | 28(4): 288-294 | No exclusion of arthropods/invertebrates; decomposition/C/N not measured | WOS |
| 150 | Takigahira and Tamawo | 2019 | Competitive responses based on kin-discrimination underlie variations in leaf functional traits in Japanese beech (Fagus crenata) seedlings | Evolutionary Ecology | 33(4): 521-531 | Invertebrates not studied | WOS |
| 151 | Diesburg et al. | 2019 | Changes in benthic invertebrate communities of central Appalachian streams attributed to hemlock woody adelgid invasion | Aquatic Sciences | 81(1): 11 | Aquatic system | WOS |
| 152 | Ruiz-Guerra et al. | 2019 | Invasive Species Appear to Disrupt the Top-Down Control of Herbivory on a Mexican Oceanic Island | Pacific Science | 73(1): 1-16 | No exclusion of arthropods/invertebrates; decomposition/C/N not measured | WOS |
| 153 | Barker et al. | 2019 | Independent and interactive effects of plant genotype and environment on plant traits and insect herbivore performance: A meta-analysis with Salicaceae | Functional Ecology | 33(3): 422-435 | Review/meta-analysis | WOS |
| 154 | Wang et al. | 2019 | Diversifying livestock promotes multidiversity and multifunctionality in managed grasslands | PNAS | 116(13): 201807354 | Decomposition/C/N not measured; effects of invertebrates not studied | WOS |
| 155 | Mahon et al. | 2019 | Experimental effects of white-tailed deer and an invasive shrub on forest ant communities | Oecologia | 191(3): 633–644 | Decomposition/C/N not measured; effects of invertebrates not studied | WOS |
| 156 | Mellado et al. | 2019 | Hemiparasites drive heterogeneity in litter arthropods: implications for woodland insectivorous birds | Austral Ecology | 44(5): 777-785 | Decomposition/C/N not measured | WOS |
| 157 | Wise and Lensing | 2019 | Impacts of rainfall extremes predicted by climate-change models on major trophic groups in the leaf litter arthropod community | Journal of Animal Ecology | 88(10): 1486-1497 | Decomposition/C/N not measured | WOS |
| 158 | Roeland et al. | 2019 | Towards an integrative approach to evaluate the environmental ecosystem services provided by urban forest | Journal of Forestry Research | 30(6): 1981–1996 | Review/meta-analysis | WOS |
| 159 | Rozanova et al. | 2019 | Arthropod rain in a temperate forest: Intensity and composition | Pedobiologia | 75: 52-56 | No exclusion of arthropods/invertebrates; decomposition/C/N not measured | WOS |
| 160 | Zhang et al. | 2019 | Invasive plants differentially affect soil biota through litter and rhizosphere pathways: a meta-analysis | Ecology Letters | 22(1): 200-210 | Review/meta-analysis | WOS |
| 161 | Entrekin et al. | 2019 | Multiple riparian-stream connections are predicted to change in response to salinization | 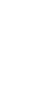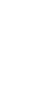 Philosophical Transactions of the Royal Society B: Biological Sciences | 374(1764): 20180042 | Review/meta-analysis | WOS |
| 162 | Ruiz-Lupion et al. | 2019 | New litter trap devices outperform pitfall traps for studying arthropod activity | Insects | 10(5): 147 | Methods paper | WOS |
| 163 | Wu et al. | 2019 | Polyploidy in invasive Solidago candensis increased plant nitrogen uptake, and abundance and activity of microbes and nematodes in soil | Soil Biology and Biochemistry | 138: 107594 | Studied impacts of an invasive plant | WOS |
| 164 | Neher and Barbercheck | 2019 | Soil microarthropods and soil health: intersection of decomposition and pest suppression in agroecosystems | Insects | 10(12): 414 | Review/meta-analysis | WOS |
| 165 | Potapov et al. | 2019 | Uncovering trophic positions and food resources of soil animals using bulk natural stable isotope composition | Biological Reviews | 94(1): 37-59 | Review/meta-analysis | WOS |
| 166 | De Smedt et al. | 2019 | Strength of forest edge effects on litter-dwelling macro-arthropods across Europe is influenced by forest age and edge properties | Diversity and Distributions | 25(6): 963-974 | Decomposition/C/N not measured | WOS |
| 167 | Tao et al. | 2019 | Soil Mesofauna Respond to the Upward Expansion of Deyeuxia purpurea in the Alpine Tundra of the Changbai Mountains, China | Plants (Basel) | 8(12): 615 | No Decomposition/C/N measured and no invertebrate exclusion | WOS |
| 168 | Bahrndorff et al. | 2018 | Diversity and metabolic potential of the microbiota associated with a soil arthropod | Scientific Reports | 8(1): 2491 | Decomposition/C/N not measured | WOS |
| 169 | Xiao et al. | 2018 | Earthworms affect plant growth and resistance against herbivores: a meta-analysis | Functional Ecology | 32(1): 150-160 | Review/meta-analysis | WOS |
| 170 | Hayes et al. | 2018 | Evidence-based logic chains demonstrate multiple impacts of trace metals on ecosystem services | Journal of Environmental Management | 223: 150-164 | Methods paper | WOS |
| 171 | Cuevas-Reyes et al. | 2018 | Effects of ferric soils on arthropod abundance and herbivory on Tibouchina heteromalla (Melastomataceae): is fluctuating asymmetry a good indicator of environmental stress? | Plant Ecology | 219(1) 69-78 | No exclusion of invertbrates | WOS |
| 172 | Walter | 2018 | Effects of changes in soil moisture and precipitation patterns on plant-mediated biotic interactions in terrestrial ecosystems | Plant Ecology | 219(12): 1449–1462 | Review/meta-analysis | WOS |
| 173 | Brueckner et al. | 2018 | Body size structure of oribatid mite communities in different microhabitats | International Journal of Acarology | 44(8): 367-373 | Decomposition/C/N not measured | WOS |
| 174 | Thunes et al. | 2018 | The red wood ant Formica aquilonia (Hymenoptera : Formicidae) may affect both local species richness and composition at multiple trophic levels in a boreal forest ecosystem | Annales Zoologici Fennici | 55(4-6): 159-172 | Decomposition/C/N not measured | WOS |
| 175 | Eeva et al. | 2018 | Leaves, berries and herbivorous larvae of bilberry Vaccinium myrtillus as sources of metals in food chains at a Cu-Ni smelter site | Chemosphere | 210: 859-866 | Decomposition/C/N not measured | WOS |
| 176 | Dossa et al. | 2018 | The cover uncovered: Bark control over wood decomposition | Journal of Ecology | 106(6): 2147-2160 | Review/meta-analysis | WOS |
| 177 | Verschut and Hamback | 2018 | A random survival forest illustrates the importance of natural enemies compared to host plant quality on leaf beetle survival rates | BMC Ecology | 18(1): 33 | Decomposition/C/N not measured | WOS |
| 178 | Schmidt | 2018 | Amber inclusions from New Zealand | Gondwana Research | 56: 135-146 | Decomposition/C/N not measured; effects of invertebrates not studied | WOS |
| 179 | Bachelot et al. | 2018 | Associations among arbuscular mycorrhizal fungi and seedlings are predicted to change with tree successional status | Ecology | 99(3): 607-620 | Decomposition/C/N not measured; effects of invertebrates not studied |  |
| 180 | Baudrot et al. | 2018 | Effects of contaminants and trophic cascade regulation on food chain stability: Application to cadmium soil pollution on small mammals - Raptor systems | Ecological Modeling | 382: 33-42 | Modeling paper | WOS |
| 181 | Jimenez-Chacon | 2018 | Fine scale determinants of soil litter fauna on a mediterranean mixed oak forest invaded by the exotic soil-borne pathogen Phytophthora cinnamomi | Forests | 9(4): 218 | Decomposition/C/N not measured | WOS |
| 182 | Pellegrino et al. | 2018 | Impact of genetically engineered maize on agronomic, environmental and toxicological traits: a meta-analysis of 21 years of field data | Scientiifc Reports | 8(1): 3113 | Review/meta-analysis | WOS |
| 183 | Hall et al. | 2018 | Invasion of Hawaiian rainforests by an introduced amphibian predator and N2-fixing tree increases soil N2O emissions | Ecosphere | 9(9): e02416 | Impacts of an amphibian; no invertebrates studied | WOS |
| 184 | Carteni et al. | 2018 | Linking plant phytochemistry to soil processes and functions: the usefulness of C-13 NMR spectroscopy | Phytochemistry Reviews | 17(5): 815–832 | Review/meta-analysis | WOS |
| 185 | Siders et al. | 2018 | Litter identity affects assimilation of carbon and nitrogen by a shredding caddisfly | Ecosphere | 9(7): e02340 | Aquatic system | WOS |
| 186 | McCary et al. | 2018 | Covariation between local and landscape factors influences the structure of ground-active arthropod communities in fragmented metropolitan woodlands | Landscape Ecology | 33(2): 225–239 | Decomposition/C/N not measured; no exclusion of invertebrates |  |
| 187 | Menta et al. | 2018 | Microarthropods biodiversity in natural, seminatural, and cultivated soils-QBS-ar approach | Applied Soil Ecology | 123: 740-743 | Methods paper | WOS |
| 188 | Vilardo et al. | 2018 | Soil arthropod composition differs between old-fields dominated by exotic plant species and remnant native grasslands | Acta Oecologica | 91: 57-64 | Decomposition/C/N not measured | WOS |
| 189 | Pausas et al. | 2018 | Fire benefits flower beetles in a Mediterranean ecosystem | PLoS ONE | 13(6): e0198951 | No exclusion of arthropods/invertebrates; decomposition/C/N not measured | WOS |
| 190 | Howard et al. | 2018 | Eco-evolutionary processes affecting plant-herbivore interactions during early community succession | Oecologia | 187(2): 547–559 | Decomposition/C/N not measured | WOS |
| 191 | De Smedt et al. | 2018 | Woodlice of Belgium: an annotated checklist and bibliography (Isopoda, Oniscidea) | ZooKeys | 801(3): 265-304 | Decomposition/C/N not measured | WOS |
| 192 | Steinwandter et al | 2018 | Structural and functional characteristics of high alpine soil macro- invertebrate communities | European Journal of Soil Biology | 86(March-April 2018): 72-80 | Decomposition/C/N not measured | WOS |
| 193 | Miao et al. | 2018 | Linkages of plant-soil interface habitat and grasshopper occurrence of typical grassland ecosystem | Ecological Indicators | 90: 324-333 | No exclusion of invertbrates | WOS |
| 194 | Cook et al. | 2018 | Laboratory Evaluation of the Direct Impact of Biochar on Adult Survival of Four Forest Insect Species | Northwest Science | 92(1): 1-8 | No exclusion of invertbrates | WOS |
| 195 | Mercado-Blanco et al. | 2018 | Belowground Microbiota and the Health of Tree Crops | Frontiers in Microbiology | 9: 1006 | Review/meta-analysis | WOS |
| 196 | Bignell | 2018 | Wood-feeding termites | Zoological Monographs | 1: 339-373 | Review/meta-analysis | WOS |
| 197 | Winckler et al. | 2017 | Benthic macroinvertebrates and degradation of phytomass as indicators of ecosystem functions in flooded rice cropping | Pesquisa Agropecuária Brasileira | 52(4): 261-270 | Aquatic system | WOS |
| 198 | Jefferson and Walker | 2017 | Biological Flora of the British Isles: Serratula tinctoria | Journal of Ecology | 105(5): 1438-1458 | Review/meta-analysis | WOS |
| 199 | Servin et al. | 2017 | Weathering in soil increases nanoparticle CuO bioaccumulation within a terrestrial food chain | Nanotoxicology | 11(1): 98-111 | No exclusion of invertbrates | WOS |
| 200 | La Riva et al. | 2017 | Arthropod communities in a selenium-contaminated habitat with a focus on ant species | Environmental Pollution | 220(A): 234-241 | No exclusion of invertbrates | WOS |
| 201 | Milano et al. | 2017 | Collembolan biodiversity in Mediterranean urban parks: impact of history, urbanization, management and soil characteristics | Applied Soil Ecology | 119: 428–437 | Decomposition/C/N not measured | WOS |
| 202 | Traunspurger et al. | 2017 | Diversity and distribution of soil micro-invertebrates across an altitudinal gradient in a tropical montane rainforest of Ecuador, with focus on free-living nematodes | Pedobiologia | 62: 28-35 | Decomposition/C/N not measured | WOS |
| 203 | Castro-Diez and Alonso | 2017 | Effects of non-native riparian plants in riparian and fluvial ecosystems: a review for the Iberian Peninsula | Limnetica | 36(2): 525-541 | Review/meta-analysis | WOS |
| 204 | Katagiri and Nana | 2017 | Effects of sika deer browsing on soil mesofauna in a thinned Japanese cypress plantation | Journal of Forest Research | 22(3): 169-176 | Decomposition/C/N not measured | WOS |
| 205 | Wells et al. | 2017 | Habitat modification by the leaf-cutter ant, Atta cephalotes, and patterns of leaf-litter arthropod communities | Environmental Entomology | 46(6): 1264–1274 | Decomposition/C/N not measured | WOS |
| 206 | Ellis and Leroux | 2017 | Moose directly slow plant regeneration but have limited indirect effects on soil stoichiometry and litter decomposition rates in disturbed maritime boreal forests | Functional Ecology | 31(3): 790-801 | No invertebrates studied | WOS |
| 207 | Cortois et al. | 2017 | Possible mechanisms underlying abundance and diversity responses of nematode communities to plant diversity | Ecosphere | 8(5): e01719 | Decomposition/C/N not measured | WOS |
| 208 | Arnold et al. | 2017 | Post-fire recovery of litter detritivores is limited by distance from burn edge | Austral Ecology | 42(1): 94-102 | Decomposition/C/N not measured | WOS |
| 209 | Gajda et al. | 2017 | Food preferences of enchytraeids | Pedobiologia | 63: 19-36 | Review/meta-analysis | WOS |
| 210 | Cole and Sheldon | 2017 | The shifting phenological landscape: Within- and between-species variation in leaf emergence in a mixed-deciduous woodland | Ecology and Evolution | 7(4): 1135-1147 | Invertebrates not studies | WOS |
| 211 | Andriuzzi and Wall | 2017 | Responses of belowground communities to large aboveground herbivores: Meta-analysis reveals biome-dependent patterns and critical research gaps | Global Change Biology | 23(9): 3857-3868 | Review/meta-analysis | WOS |
| 212 | Tsunodo and van Dam | 2017 | Root chemical traits and their roles in belowground biotic interactions | Pedobiologia | 65: 58-67 | Review/meta-analysis | WOS |
| 213 | Bayliss et al. | 2017 | Testing genotypic variation of an invasive plant species in response to soil disturbance and herbivory | Oecologia | 183(4): 1135–1141 | No exclusion of invertbrates | WOS |
| 214 | Dombos | 2017 | EDAPHOLOG monitoring system: automatic, real-time detection of soil microarthropods | Methods in Ecology and Evolution | 8(3): 313-321 | Methods paper | WOS |
| 215 | Gonzalez and Lodge | 2017 | Soil biology research across latitude, elevation and disturbance gradients: a review of forest studies from Puerto Rice during the past 25 years | Forests | 8(6): 178 | Review/meta-analysis | WOS |
| 216 | Costa-Milaneza et al. | 2017 | Influence of soil granulometry on average body size in soil ant assemblages: implications for bioindication | Perspectives in Ecology and Conservation | 15(2): 102-108 | Review/meta-analysis | WOS |
| 217 | Thoresen et al. | 2017 | Invasive rodents have multiple indirect effects on seabird island invertebrate food web structure | Ecological Applications | 27(4): 1190-1198 | No exclusion of invertbrates | WOS |
| 218 | Stephan et al. | 2017 | Long-term deer exclosure alters soil properties, plant traits, understory plant community and insect herbivory, but not the functional relationships among them | Oecologia | 184(3): 685–699 | No exclusion of invertbrates | WOS |
| 219 | Eldridge et al. | 2017 | Direct and Indirect Effects of Herbivore Activity on Australian Vegetation | Ecology and Evolution | 10(1): 557-568 | Review/meta-analysis | WOS |
| 220 | Qin et al. | 2017 | Wood ash application increases pH but does not harm the soil mesofauna | Environmental Pollution | 224: 581-589 | No exclusion of invertbrates | WOS |
| 221 | Adouzi et al. | 2017 | Detecting pyrethroid resistance in predatory mites inhabiting soil and litter: an in vitro test | Pest Management Science | 73(6): 1258-1266 | No exclusion of invertbrates | WOS |
| 222 | Klarner et al. | 2017 | Trophic niches, diversity and community composition of invertebrate top predators (Chilopoda) as affected by conversion of tropical lowland rainforest in Sumatra (Indonesia) | PLoS ONE | 12(8): e0180915 | No exclusion of invertbrates | WOS |
| 223 | Gan and Wickings | 2017 | Soil ecological responses to pest management in golf turf vary with management intensity, pesticide identity, and application program | Agriculture, Ecosystems & Environment | 246: 66-77 | Decomposition/C/N not measured | WOS |
| 224 | Gornostaeva et al. | 2017 | The effect of fluorinated compounds on living organisms (review) | Theoretical and Applied Ecology | 2017(1): 14–24 | Review/meta-analysis | WOS |
| 225 | De Smedt et al. | 2016 | Complementary distribution patterns of arthropod detritivores (woodlice and millipedes) along forest edge-to-interior gradients | Insect Conservation and Diversity | 9(5): 456-469 | Decomposition/C/N not measured | WOS |
| 226 | Fuhrer et al. | 2016 | Current and future ozone risks to global terrestrial biodiversity and ecosystem processes | Ecology and Evolution | 6(24): 8785-8799 | Review/meta-analysis | WOS |
| 227 | Menz et al. | 2016 | Dispersal-limited detritivores in fire-prone environments: persistence and populaiton structure of terrestrial amphipods (Talitridae) | International Journal of Wildland Fire | 25(7): 753-761 | Decomposition/C/N not measured | WOS |
| 228 | Natusch et al. | 2016 | Communally Nesting Migratory Birds Create Ecological Hot-Spots in Tropical Australia | PLoS ONE | 11(10): e0162651. | No exclusion of invertbrates | WOS |
| 229 | Nardi et al. | 2016 | Compartmentalization of microbial communities that inhabit the hindguts of millipedes | Arthropod Structure & Development | 45(5): 462-474 | No exclusion of invertbrates | WOS |
| 230 | Minor et al. | 2016 | Effects of cushion plants on high-altitude soil microarthropod communities: cushions increase abundance and diversity of mites (Acari), but not springtails (Collembola) | Arctic, Antarctic, and Alpine Research | 48: 485-500 | Decomposition/C/N not measured | WOS |
| 231 | Toth et al. | 2016 | Effects of set-aside management on soil macrodecomposers in Hungary | Applied Soil Ecology | 99: 89-97 | Decomposition/C/N not measured | WOS |
| 232 | Cameron et al. | 2016 | Global meta-analysis of the impacts of terrestrial invertebrate invaders on species, communities, and ecosystems | Global Ecology and Biogeography | 25(5): 596-606 | A meta-analysis | WOS |
| 233 | Schowalter | 2016 | Herbivory | Insect Ecology: An Ecosystem Approach (4th ed) (Book) | 406-442 | Review/meta-analysis | WOS |
| 234 | Gongalsky et al. | 2016 | Diversity of the soil biota in burned areas of southern taiga forests (Tver oblast) | Eurasian Soil Science | 49(3): 358–366 | No exclusion of invertbrates | WOS |
| 235 | Maceda-Veiga | 2016 | Impacts of the invader giant reed (Arundao donax) on riparian habitats and ground arthropod communities | Biological Invasions | 18(3): 731–749 | Impacts of an invasive plant | WOS |
| 236 | Schmidt | 2016 | Photoautotrophic microorganisms as a carbon source for temperate soil invertebrates | Biology Letters | 12(1): 20150646 | Decomposition/C/N not measured | WOS |
| 237 | Filipiak | 2016 | Pollen stoichiometry may influence detrital terrestrial and aquatic food webs | Frontiers in Ecology and Evolution | 4: 138 | Decomposition/C/N not measured | WOS |
| 238 | Chertov | 2016 | Quantification of soil fauna metabolites and dead mass as humification sources in forest soils | Eurasian Soil Science | 49(1): 77-88 | Decomposition/C/N not measured | WOS |
| 239 | Schaller et al. | 2016 | Reed litter Si content affects microbial community structure and lipid composition of an invertebrate shredder during aquatic decomposition | Limnologica | 57: 14-22 | Aquatic system | WOS |
| 240 | Maute et al. | 2016 | Effects of two locust control methods on wood-eating termites in arid Australia | Journal of Insect Conservation | 20(1): 107–118 | No exlusion of invertebrates | WOS |
| 241 | Santos et al. | 2016 | Characterization of soil macrofauna in grain production systems in the Southeastern state of Piaui, Brazil | Pesquisa Agropecuária Brasileira | 51(9): 1466-1475 | Decompositin/C/N not measured | WOS |
| 242 | Birkhofer | 2016 | Regional conditions and land-use alter the potential contribution of soil arthropods to ecosystem services in grasslands | Frontiers in Ecology and Evolution | 3: 150 | Decomposition/C/N not measured | WOS |
| 243 | Goncharov et al. | 2016 | Short-term incorporation of freshly fixed plant carbon into the soil animal food web: field study in a spruce forest | Ecological Research | 31(6): 923-933 | Decomposition/C/N not measured | WOS |
| 244 | Potapov and Tiunov | 2016 | Stable isotope composition of mycophagous collembolans versus mycotrophic plants: do soil invertebrates feed on mycorrhizal fungi? | Soil Biology and Biochemistry | 93: 115-118 | Decomposition/C/N not measured; no invertebrate exclusion | WOS |
| 245 | Duyar and Makineci | 2016 | The seasonal variation of arthropods living on forest soil at different altitudes in fir (Abies nordmanniana subsp. Bornmulleriana) ecosystem in Bolu-Aladag | Forestist | 66(2): 572-586 | Decomposition/C/N not measured | WOS |
| 246 | Namba and Ohdachi | 2016 | Top-down cascade effects of the long-clawed shrew (Sorex unguiculatus) on the soil invertebrate community in a cool-temperate forest | Mammal Study | 41(3): 119-130 | Impacts of a shrew on litter decomposition | WOS |
| 247 | Tuovinen et al. | 2016 | Transfer of elements relevant to nuclear fuel cycle from soil to boreal plants and animals in experimental meso- and microcosms | Science of the Total Environment | 539: 252-261 | Decomposition/C/N not measured | WOS |
| 248 | Harvey et al. | 2016 | Short-term seasonal habitat facilitation mediated by an insect herbivore | Basic and Applied Ecology | 17(5): 447-454 | Decomposition/C/N not measured; no invertebrate exclusion | WOS |
| 249 | Gregory et al. | 2016 | Agroecological and social characteristics of New York city community gardens: contributions to urban food security, ecosystem services, and environmental education | Urban Ecosystems | 19(2): 763–794 | Decomposition/C/N not measured | WOS |
| 250 | La Pierre and Smith | 2016 | Soil nutrient additions increase invertebrate herbivore abundances, but not herbivory, across three grassland systems | Oecologia | 180(2): 485–497 | No exlusion of invertebrates | WOS |
| 251 | Kaminski et al. | 2016 | Effects of chemical elements in the trophic levels of natural salt marshes | Environmental Geochemistry and Health | 38(3): 783–810 | No exclusion of invertbrates | WOS |
| 252 | Schowalter | 2016 | Community structure | Insect Ecology: An Ecosystem Approach (4th ed) (Book) | 292-324 | Review/meta-analysis | WOS |
| 253 | Greenstone | 2016 | Sampling Epigeal Arthropods: A Permanent, Sheltered, Closeable Pitfall Trapping Station | Journal of Entomological Service | 51(1): 87-93 | Methods paper | WOS |
| 254 | Bowie et al. | 2016 | Persistence of biodiversity in a dryland remnant within an intensified dairy farm landscape | New Zealand Journal of Ecology | 40(1): 121-130 | No exclusion of invertbrates | WOS |
| 255 | Mueller et al. | 2016 | Light, earthworms, and soil resources as predictors of diversity of 10 soil invertebrate groups across monocultures of 14 tree species | Soil Biology and Biochemistry | 92: 184-198 | No exclusion of invertbrates | WOS |
| 256 | King | 2016 | Where do eusocial insects fit into soil food webs? | Soil Biology and Biochemistry | 102: 55-62 | Review/meta-analysis | WOS |
| 257 | Van Stan et al. | 2015 | A review and evaluation of forest canopy epiphyte roles in the partitioning and chemical alteratoin of precipitation | Science of the Total Environment | 536: 813-824 | Review/meta-analysis | WOS |
| 258 | Sun et al. | 2015 | Across-trophic variation of potassium, calcium, and magnesium stoichiometric traits in a parasitism food chain across temperate and subtropical biomes | Crop and Pasture Science | 66(12): 1290-1297 | Decomposition/C/N not measured | WOS |
| 259 | Tiunov et al. | 2015 | Stable isotope composition (delta C-13 and delta N-15 values) of slime molds: placing bacterivorous soil protozoans in the food web context | Rapid Communications in Mass Spectrometry | 29(16): 1465-1472 | No exlusion of invertebrates | WOS |
| 260 | Milcu et al. | 2015 | Aphid honeydew-induced changes in soil biota can cascade up to tree crown architecture | Pedobiologia | 58(4): 119-127 | Impacts of honeydew on decomposition; not invertebrate-induced | WOS |
| 261 | Xu et al. | 2015 | Context dependency of the density-body mass relationship in litter invertebrates along an elevational gradient | Soil Biology and Biochemistry | 88: 323-332 | Decomposition/C/N not measured | WOS |
| 262 | Bourguignon et al. | 2015 | Influence of Soil Properties on Soldierless Termite Distribution | PLoS ONE | 10(8): e0135341 | No exlusion of invertebrates | WOS |
| 263 | Liu et al. | 2015 | Effectiveness of afforested shrub plantation on ground-active arthropod communities and trophic structure in desertified regions | CATENA | 125: 1-9 | Decomposition/C/N not measured | WOS |
| 264 | Kolb et al. | 2015 | Effects of nesting cormorants (Phalacrocorax carbo) on soil chemistry, microbial communities, and soil fauna | Ecosystems | 18(4): 643–657 | Impacts of a seabird | WOS |
| 265 | McCary et al. | 2015 | Effects of woodland restoration and management on the community of surface-active arthropods in the metropolitan Chicago region | Biological Conservation | 190: 154-166 | Decomposition/C/N not measured | WOS |
| 266 | Callejas-Chavero et al. | 2015 | Soil Microarthropods and Their Relationship to Higher Trophic Levels in the Pedregal de San Angel Ecological Reserve, Mexico | Journal of Insect Science | 15(1): 59 | Decomposition/C/N not measured | WOS |
| 267 | Mori et al. | 2015 | Concordance and discordance between taxonomic and functional homogenization: responses of soil mite assemblages to forest conversion | Oecologia | 179(2): 527-35 | No exclusion of invertbrates | WOS |
| 268 | Hulber et al. | 2015 | Insect herbivory in alpine grasslands is constrained by community and host traits | Journal of Vegetation Science | 26(4): 663-673 | Decomposition/C/N not measured | WOS |
| 269 | Hyodo et al. | 2015 | Dependence of diverse consumers on detritus in a tropical rain forest food web as revealed by radiocarbon analysis | Functional Ecology | 29(3): 423-429 | Decomposition/C/N not measured | WOS |
| 270 | Mori et al. | 2015 | Biotic homogenization and differentiation of soil faunal communities in the production forest landscape: taxonomic and functional perspectives | Oecologia | 177: 533–544 | Decomposition/C/N not measured | WOS |
| 271 | Fluery et al. | 2015 | Seedling fate across different habitats: The effects of herbivory and soil fertility | Basic and Applied Ecology | 16(2): 141-151 | Decomposition/C/N not measured | WOS |
| 272 | Doblas-Miranda and Work | 2015 | Localized Effects of Coarse Woody Material on Soil Oribatid Communities Diminish over 700 Years of Stand Development in Black-Spruce-Feathermoss Forests | Forests | 6(4): 914-928 | Decomposition/C/N not measured | WOS |
| 273 | Ladin et al. | 2015 | Is brood parasitism related to host nestling diet and nutrition? | The Auk | 132(3): 717–734 | Decomposition/C/N not measured | WOS |
| 274 | Flacher et al. | 2015 | Competition with wind-pollinated plant species alters floral traits of insect-pollinated plant species | Scientific Reports | 5(1): 13354 | Decomposition/C/N not measured | WOS |
| 275 | Kirchberger et al. | 2015 | Experimental evaluation of herbivory on live plant seedlings by the earthworm Lumbricus terrestris L. in the presence and absence of soil surface litter | PLoS ONE | 10(4): e0123465 | Decomposition/C/N not measured | WOS |
| 276 | Mortard et al. | 2015 | How invasion by Ailanthus altissima transforms soil and litter communities in a temperate forest ecosystem | Biological Invasions | 17(6): 1817–1832 | Impacts of an invasive plant | WOS |
| 277 | Brygadyrenko | 2015 | Influence of moisture conditions and mineralization of soil solution on structure of litter macrofauna of the deciduous forests of Ukraine steppe zone | Biosystems Diversity | 23(1): 50-65 | Decomposition/C/N not measured | WOS |
| 278 | Pinkalski | 2015 | Quantification of ant manure deposition in a tropical agroecosystem: implications for host plant nitrogen acquisition | Ecosystems | 18(8): 1373–1382 | Decomposition/C/N not measured | WOS |
| 279 | Mulder et al. | 2015 | Leaf damage by herbivores and pathogens on New Zealand islands that differ in seabird densities | New Zealand Journal of Ecology | 39(2): 1-10 | No exclusion of invertbrates | WOS |
| 280 | Scheunemann et al. | 2015 | Roots rather than shoot residues drive soil arthropod communities of arable fields | Oecologia | 179: 1135–1145 | Decomposition/C/N not measured | WOS |
| 281 | Carron et al. | 2015 | Spatial heterogeneity of soil quality around mature oil palms receiving mineral fertilization | European Journal of Soil Biology | 66: 24-31 | Invertebrates not studied; decomposition/C/N not measured | WOS |
| 282 | Stevens et al. | 2015 | Root Chemistry in Populus tremuloides: Effects of Soil Nutrients, Defoliation, and Genotype | Journal of Chemical Ecology | 40(1): 31–38 | Invertebrates not studied | WOS |
| 283 | Wu et al. | 2014 | The brown-world role of insectivores: Frogs reduce plant growth by suppressing detritivores in an alpine meadow | Basic and Applied Ecology | 15(1): 66-74 | No invertebrate exclusion | WOS |
| 284 | Walker et al. | 2014 | A metagenomics-based approach to the top-down effect on the detritivore food web: a salamanders influence on fungal communities within a deciduous forest | Ecology and Evolution | 4(21): 4106–4116 | Decomposition/C/N not measured; salamander effects | WOS |
| 285 | Santorufo et al. | 2014 | An assessment of the influence of the urban environment on collembolan communities in soils using taxonomy- and trait-based approaches | Applied Soil Ecology | 78: 48-56 | Decomposition/C/N not measured | WOS |
| 286 | Joyce | 2014 | Ecological consequences and restoration potential of abandoned wet grasslands | Ecological Engineering | 66: 91-102 | Review/meta-analysis | WOS |
| 287 | Chen et al. | 2014 | Aromatic plants play an important role in promoting soil biological activity related to nitrogen cycling in an orchard ecosystem | Science of the Total Environment | 472: 939-946 | Invertebrates not studied | WOS |
| 288 | Book et al. | 2014 | Cellulolytic Streptomyces Strains Associated with Herbivorous Insects Share a Phylogenetically Linked Capacity To Degrade Lignocellulose | Applied and Environmental Microbiology | 80(15): 4692–4701 | Decomposition/C/N not measured | WOS |
| 289 | Low et al. | 2014 | Elevated volatile concentrations in high-nutrient plants: do insect herbivores pay a high price for good food? | Ecological Entomology | 39(4): 480-91 | Decomposition/C/N not measured | WOS |
| 290 | Maraun et al. | 2014 | Changes in the community composition and trophic structure of microarthropods in sporocarps of the wood decaying fungus Fomitopsis pinicola along an altitudinal gradient | Applied Soil Ecology | 84: 16–23 | Decomposition/C/N not measured | WOS |
| 291 | Kutovaya et al. | 2014 | Communities of microorganisms and invertebrates in soil-like bodies of soccer fields in Moscow oblast | Eurasian Soil Science | 47(11): 1107–1115 | Decomposition/C/N not measured | WOS |
| 292 | Crotty et al. | 2014 | Divergence of feeding channels within the soil food web determined by ecosystem type | Ecology and Evolution | 4(1): 1-13 | Decomposition/C/N not measured | WOS |
| 293 | Bassett | 2014 | Impacts on invertebrate fungivores: a predictable consequence of ground-cover weed invasion? | Biodiversity and Conservation | 23(4): 791-810 | Decomposition/C/N not measured | WOS |
| 294 | Alatalo and Little | 2014 | Simulated global change: contrasting short and medium term growth and reproductive responses of a common alpine/Arctic cushion plant to experimental warming and nutrient enhancement | SpringerPlus | 3: 157 | No invertebrates studied | WOS |
| 295 | Ribeiro et al. | 2014 | Influence of mineral fertilization on edaphic fauna in Acacia auriculiformis (A. Cunn) plantations | Revista Brasileira de Ciência do Solo | 38(1): 39-49 | Decomposition/C/N not measured | WOS |
| 296 | Yang and Gratton | 2014 | Insects as drivers of ecosystem processes | Current Opinion in Insect Science | 2: 26-32 | Review/meta-analysis | WOS |
| 297 | Raub et al. | 2014 | No bottom-up effects of food addition on predators in a tropical forest | Basic and Applied Ecology | 15: 59-65 | Decomposition/C/N not measured | WOS |
| 298 | Vaz et al. | 2014 | Effects of burn status and conditioning on colonization of wood by stream macroinvertebrates | Freshwater Science | 33(3): 832-846 | Aquatic system | WOS |
| 299 | A'Bear et al. | 2014 | Putting the 'upstairs-downstairs' into ecosystem service: what can aboveground-belowground ecology tell us? | Biological Control | 75: 97-107 | Review/meta-analysis | WOS |
| 300 | A'Bear et al. | 2014 | Size matters: what have we learnt from microcosm studies of decomposer fungus-invertebrate interactions? | Soil Biology and Biochemistry | 78: 274-283 | Review/meta-analysis | WOS |
| 301 | Bokhorst and Wardle | 2014 | Snow fungi as a food source for micro-arthropods | European Journal of Soil Biology | 60: 77-80 | Decomposition/C/N not measured | WOS |
| 302 | Best and Welsh | 2014 | The trophic role of a forest salamander: impacts on invertebrates, leaf litter retention, and the humification process | Ecosphere | 5(2): 1-19 | Impacts of salamanders; decomposition/C/N not measured | WOS |
| 303 | Kuebbing et al. | 2014 | Two co-occurring invasive woody shrubs alter soil properties and promote subdominant invasive species | Journal of Applied Ecology | 51(1): 124-133 | No invertebrate exclusion | WOS |
| 304 | Ferlian and Scheu | 2014 | Shifts in trophic interactions with forest type in soil generalist predators as indicated by complementary analyses of fatty acids and stable isotopes | Oikos | 123(10): 1182-1191 | No invertebrate exclusion | WOS |
| 305 | Lemanski et al. | 2014 | Fertilizer addition lessens the flux of microbial carbon to higher trophic levels in soil food webs of grassland | Oecologia | 176(2): 487-496 | No invertebrate exclusion | WOS |
| 306 | Klarner et al. | 2014 | Trophic shift of soil animal species with forest type as indicated by stable isotope analysis | Oikos | 123(10): 1173-1181 | No invertebrate exclusion | WOS |
| 307 | Cierjacks et al. | 2013 | Biological flora of the British Isles: Robinia pseudoacacia | Journal of Ecology | 101(6): 1623-1640 | Review/meta-analysis | WOS |
| 308 | Lavorel | 2013 | Plant functional effects on ecosystem services | Journal of Ecology | 101(1): 4-8 | Review/meta-analysis | WOS |
| 309 | Mulder et al. | 2013 | Connecting the green and brown worlds: allometric and stoichiometric predictability of above- and belowground networks | Advances in Ecological Research | 49: 69-175 | Review/meta-analysis | WOS |
| 310 | Diaz-Anguilar et al. | 2013 | Influence of stand composition on predatory mite (Mesostigmata) assemblages from the forest floor in western Canadian boreal mixedwood forests | Forest Ecology and Management | 309: 105-114 | No invertebrate exclusion | WOS |
| 311 | Souza-Silva et al. | 2013 | Food resource availability in a quartzite cave in the Brazilian Montane Atlantic Forest | Journal of Cave and Karst Studies | 75(3): 177–188 | Decomposition/C/N not measured | WOS |
| 312 | Pregitzer et al. | 2013 | Genetic by environmental interactions affect plant-soil linkages | Ecology and Evolution | 3(7): 2322-2333 | No invertebrate exclusion | WOS |
| 313 | Pantankar et al. | 2013 | Permafrost-driven differences in habitat quality determine plant response to gall-inducing mite herbivory | Journal of Ecology | 101(4): 1042-1052 | Decomposition/C/N not measured | WOS |
| 314 | Griffith et al. | 2013 | Herbivore behavior in the anecic earthworm species Lumbricus terrestris L. | European Journal of Soil Biology | 55: 62-65 | Decompositon/C/N not measured | WOS |
| 315 | Welsh et al. | 2013 | Woodland salamanders as metrics of forest ecosystem recovery: a case study from California's redwoods | Ecosphere | 4(5): 1-25 | Invertebrates not studied | WOS |
| 316 | Gorman et al. | 2013 | Species identity influences belowground arthropod assemblages via functional traits | AoB Plants | 5: plt049 | No exclusion of invertbrates | WOS |
| 317 | Zaller et al. | 2013 | Herbivory of an invasive slug is affected by earthworms and the composition of plant communities | BMC Ecology | 13(1): 20 | Decompositon/C/N not measured | WOS |
| 318 | Krab et al. | 2013 | How extreme is an extreme climatic event to a subarctic peatland springtail community? | Soil Biology and Biochemistry | 59: 6-24 | Decomposition/C/N not measured | WOS |
| 319 | Almeida Pereira et al. | 2013 | Litter decomposition, diversity and functionality of soil invertebrates in an atlantic rain forest fragment | Bioscience Journal | 29(5): 1316-1326 | Review/meta-analysis | WOS |
| 320 | Barber et al. | 2013 | Linking agricultural practices, mycorrhizal fungi, and traits mediating plant-insect interactions | Ecological Applications | 23(7): 1519-30 | Decomposition/C/N not measured | WOS |
| 321 | Colloff et al. | 2013 | Natural pest control in citrus as an ecosystem service: integrating ecology, economics, and management at the farm scale | Biological Control | 67(2): 170-177 | Review/meta-analysis | WOS |
| 322 | Wurst et al. | 2013 | Plant-mediated links between detritivores and aboveground herbivores | Frontiers in Plant Science | 4: 380 | Review/meta-analysis | WOS |
| 323 | Bastow | 2013 | Succession, resource processing, and diversity in detrital food webs | Soil Ecology and Ecosystem Services (book) | 117-135 | Review/meta-analysis | WOS |
| 324 | Bressette et al. | 2012 | Beyond the browse line: complex cascade effects mediated by white-tailed deer | Oikos | 121(11): 1749-1760 | Review/meta-analysis | WOS |
| 325 | Loiola et al. | 2012 | Underdispersion of anti-herbivore defence traits and phylogenetic structure of cerrado tree species at fine spatial scale | Journal of Vegetation Science | 23(6): 1095-1104 | No exclusion of invertbrates | WOS |
| 326 | Cooke and Leishman | 2012 | Tradeoffs between foliar silicon and carbon-based defences: evidence from vegetation communities of contrasting soil types | Oikos | 121(12): 2052-2060 | No exclusion of invertbrates | WOS |
| 327 | Eldridge et al. | 2012 | Soil Disturbance by Invertebrates in a Semi-arid Eucalypt Woodland: Effects of Grazing Exclusion, Faunal Reintroductions, Landscape and Patch Characteristics | Proceedings of the Linnean Society of New South Wales | 134: A11-A18 | Decomposition/C/N not measured | WOS |
| 328 | Cornelissen et al. | 2012 | Controls on coarse wood decay in temperate tree species: birth of the loglife experiment | AMBIO | 41(Supp. 3): 231-245 | Review/meta-analysis | WOS |
| 329 | Bokhorst et al. | 2012 | Extreme winter warming events more negatively impact small rather than large soil fauna: shift in community composition explained by traits not taxa | Global Change Biology | 18(3): 1152-1162 | Decomposition/C/N not measured | WOS |
| 330 | McCall and Pennings | 2012 | Geographic variation in salt marsh structure and function | Oecologia | 170(3): 777-787 | Decomposition/C/N not measured | WOS |
| 331 | Templer et al. | 2012 | Impact of a reduced winter snowpack on litter arthropod abundance and diversity in a northern hardwood forest ecosystem | Biology and Fertility of Soils | 48(4): 413–424 | Decomposition/C/N not measured | WOS |
| 332 | Edwards et al. | 2012 | Impacts of logging and rehabilitation on invertebrate communities in tropical rainforests of northern Borneo | Journal of Insect Conservation | 16(4): 591–599 | Decomposition/C/N not measured | WOS |
| 333 | Pozo et al. | 2012 | Nectar yeasts of two southern Spanish plants: the roles of immigration and physiological traits in community assembly | FEMS Microbiology Ecology | 80(2): 281-293 | Decomposition/C/N not measured |  |
| 334 | Melguizo-Ruiz et al. | 2012 | Potential drivers of spatial structure of leaf-ltter food webs in south-western European beech forests | Pedobiologia | 55(6): 311-319 | Decomposition/C/N not measured | WOS |
| 335 | Radea and Margarita | 2012 | Soil arthropod communities and population dynamics following wildfires in pine forests of the mediterranean basin: a review | Israel Journal of Ecology and Evolution | 58(2-3): 137-149 | A review/meta-analysis; decomposition/C/N not measured | WOS |
| 336 | Erdmann et al. | 2012 | Regional factors rather than forest type drive the community structure of soil living oribatid mites (Acari, Oribatida) | Experimental and Applied Acarology | 57(2): 157-169 | No exclusion of invertbrates | WOS |
| 337 | Arnold et al. | 2012 | Herbivory, pathogens, and epiphylls in Araucaria Forest and ecologically-managed tree monocultures | Forest Ecology and Management | 262(6): 1041-1046 | Decomposition/C/N not measured | WOS |
| 338 | Huerta and Van der Wal | 2012 | Soil macroinvertebrates' abundance and diversity in home gardens in Tabasco, Mexico, vary with soil texture, organic matter, and vegetation cover | European Journal of Soil Biology | 50: 68-75 | Decomposition/C/N not measured | WOS |
| 339 | Schon et al. | 2012 | Litter effects on seedling establishment interact with seed position and earthworm activity | Plant Biology (Stuttgart) | 14(1): 163-70 | Decomposition/C/N not measured | WOS |
| 340 | Crotty et al. | 2012 | Protozoan Pulses Unveil Their Pivotal Position Within the Soil Food Web | Microbial Ecology | 63(4): 905-918 | No exlusion of invertebrates | WOS |
| 341 | Schon et al. | 2012 | Relationship between Food Resource, Soil Physical Condition, and Invertebrates in Pastoral Soils | Soil Science Society of America Journal | 76(5): 1644-1654 | No exclusion of invertbrates | WOS |
| 342 | Crotty et al. | 2012 | Using stable isotopes to differentiate trophic feeding channels within soil food webs | Eukaryotic Microbiology | 59(6): 520-526 | Decomposition/C/N not measured | WOS |
| 343 | Crotty et al. | 2011 | Tracking the flow of bacterially derived C-13 and N-15 through soil faunal feeding channels | Rapid Communications in Mass Spectrometry | 25(11): 1503-1513 | No exclusion of arthropods | WOS |
| 344 | Pelini et al. | 2011 | Heating up the forest: open-top chamber warming manipulation of arthropod communities at Harvard and Duke Forests | Methods in Ecology and Evolution | 2(5): 534-540 | Decomposition/C/N not measured | WOS |
| 345 | Daugherty | 2011 | Host plant quality, spatial heterogeneity, and the stability of mite predator-prey dynamics | Experimental and Applied Acarology | 53(4): 311-322 | No exclusion of invertbrates | WOS |
| 346 | Ald et al. | 2011 | Reconstructing the soil food web of a 100 million-year-old forest: the case of the mid-Cretaceous fossils in the amber of Charentes (SW France) | Soil Biology and Biochemistry | 43(4): 1-10 | Review/meta-analysis | WOS |
| 347 | Montero et al. | 2011 | Seasonal assemblages of epigean arthropods in a quebracho forest (Schinopsis balansae) in the Humid Chaco | Revista Colombiana de Entomología | 37(2): 294-304 | Decomposition/C/N not measured | WOS |
| 348 | Schon et al. | 2011 | Soil fauna in sheep-grazed hill pastures under organic and conventional livestock management and in an adjacent ungrazed pasture | Pedobiologia | 54(3): 161-168 | Decomposition/C/N not measured; no invertebrate exclusion | WOS |
| 349 | Ober and DeGroote | 2011 | Effects of litter removal on arthropod communities in pine plantations | Biodiversity and Conservation | 20: 1273–1286 | No exlusion of invertebrates | WOS |
| 350 | Prevosto et al. | 2011 | Effects of different site preparation treatments on species diversity, composition, and plant traits in Pinus halepensis woodlands | Plant Ecology | 212(4): 627-638 | No exlusion of invertebrates | WOS |
| 351 | Heiner et al. | 2011 | Stable isotope N-15 and C-13 labelling of different functional groups of earthworms and their casts: A tool for studying trophic links | Pedobiologia | 54(3): 169–175 | Decomposition/C/N not measured | WOS |
| 352 | Maraun et al. | 2011 | Stable isotopes revisited: their use and limits for oribatid mite trophic ecology | Soil Biology and Biochemistry | 43(5): 877-882 | Review/meta-analysis | WOS |
| 353 | Watson | 2011 | A productivity-based explanation for woodland bird declines: poorer soils yield less food | Emu | 111(1): 10-18 | Invertebrates not studied | WOS |
| 354 | McMullan-Fisher | 2011 | Fungi and fire in Australian ecosystems: a review of current knowledge, management implications and future directions | Australian Journal of Botany | 59(1): 70-90 | Review/meta-analysis | WOS |
| 355 | Steinbauer and Peveling | 2011 | The impact of the locust control insecticide fipronil on termites and ants in two contrasting habitats in Northern Australia | Crop Protection | 30(7): 814-825 | Decomposition/C/N not measured | WOS |
| 356 | Tiley | 2010 | Biological flora of the British Isles: Cirsium arvense Scop. | Journal of Ecology | 98(4): 938-983 | Review/meta-analysis | WOS |
| 357 | Yamamoto et al. | 2010 | Detection of effects of high trophic level predator, sorex unguiculatus (Soricidae, Mammalia), on a soil microbial community in a cool temperature forest in Hokkaido, using the ARISA method | Microbes and Environments | 25(3): 197-203 | Decomposition/C/N not measured | WOS |
| 358 | Leon-Gamboa et al. | 2010 | Effect of pine planations on soil arthropods in a high Andean forest | Revista de Biologia Tropical | 58(3): 1031-48 | Decomposition/C/N not measured | WOS |
| 359 | Christiansen and Lavigne | 2010 | Effects of the 1988 fires in Yellowstone National Park, USA, on the ant populations (Hymenoptera: Formicidae) | Journal of the Entomological Research Society | 12(3): 29-37 | Studied fire effects on ant populations; decomposition/C/N not measured | WOS |
| 360 | Rodrigques Capitulo et al. | 2010 | Global changes in pampen lowland streams (Agentina): implications for biodiversity and functioning | Hydrobiologia | 657(1): 53-70 | Review/meta-analysis: aquatic system | WOS |
| 361 | Giurginca et al. | 2010 | Assessment of potentially toxic metals of concentration in karst areas of the Mehedinti Plateau Geopark (Romania) | Carpathian Journal of Earth and Environmental Science | 5(1): 103-110 | Decomposition/C/N not measured | WOS |
| 362 | Lindroth | 2010 | Impacts of elevated atmospheric CO2 on forests: phytochemistry, trophic interactions, and ecosystem dynamics | Journal of Chemical Ecology | 36(1): 2-21 | Review/meta-analysis | WOS |
| 363 | Hawke and Clark | 2010 | Isotopic signatures (13C/12C; 15N/14N) of blue penguin burrow soil invertebrates: carbon sources and trophic relationships | New Zealand Journal of Ecology | 37(4): 313-321 | Decomposition/C/N not measured; isotope study | WOS |
| 364 | Wolkovich | 2010 | Nonnative grass litter enhances grazing arthropod assemblages by increasing native shrub growth | Ecology | 91(3) :756-66 | Decomposition/C/N not measured | WOS |
| 365 | Koetsier et al. | 2010 | Present effects of past wildfires on leaf litter breakdown in stream ecosystems | Western North American Naturalist | 70(2): 164-174 | Aquatic system | WOS |
| 366 | Hagvar et al. | 2010 | Effect of simulated environmental change on alpine soil arthropods | Global Change Biology | 15(12): 2972-2980 | Decomposition/C/N not measured | WOS |
| 367 | Beaulieu et al. | 2010 | The canopy starts at 0.5m: predatory mites (acari:mesostigmata) differ between rain forest floor soil and suspended at any height | Biotropica | 42(6): 704-709 | Decomposition/C/N not measured | WOS |
| 368 | Garibaldi et al. | 2010 | Nutrient supply and bird predation additively control insect herbivory and tree growth in two contrasting forest habitats | Oikos | 119(2): 337-349 | No exclusion of invertbrates | WOS |
| 369 | Sackett et al. | 2010 | Linking soil food web structure to above- and belowground ecosystem processes: a meta-analysis | Oikos | 119(12): 1984-1992 | Review/meta-analysis | WOS |
| 370 | Magbanua et al. | 2010 | Responses of stream macroinvertebrates and ecosystem function to conventional, integrated and organic farming | Journal of Applied Ecology | 47(5): 1014-1025 | Aquatic system | WOS |
| 371 | Zvereva and Kozlov | 2010 | Responses of terrestrial arthropods to air pollution: a meta-analysis | Environmental Science and Pollution Research | 7(2): 297-311 | Review/meta-analysis | WOS |
| 372 | Bairstow et al. | 2010 | Leaf miner and plant galler species richness on Acacia: relative importance of plant traits and climate | Oecologia | 163(2): 437-448 | No exclusion of invertbrates | WOS |
| 373 | Bishop et al. | 2010 | N-P Co-Limitation of Primary Production and Response of Arthropods to N and P in Early Primary Succession on Mount St. Helens Volcano | PLoS ONE | 5(10): e13598 | No exclusion of invertbrates | WOS |
| 374 | Underwood and Quinn | 2010 | Response of ants and spiders to prescribed fire in oak woodlands of California | Journal of Insect Conservation | 14(4): 359-366 | Decomposition/C/N not measured | WOS |
| 375 | de Bello et al. | 2010 | Towards an assessment of multiple ecosystem processes and services via functional traits | Biodiversity and Conservation | 19(10): 2873–2893 | Review/meta-analysis | WOS |
| 376 | Donoso et al. | 2010 | Trees as templates for tropical litter arthropod diversity | Oecologia | 164(1): 201-211 | Decomposition/C/N not measured | WOS |
| 377 | Hawke and Clark | 2010 | Incorporation of the invasive mallow Lavatera arborea into the food web of an active seabird island | Biological Invasions | 12(6): 1805–1814 | No exlusion of invertebrates | WOS |
| 378 | Spiller et al. | 2010 | Marine subsidies have multiple effects on coastal food webs | Ecology | 91(5):1424-1434 | Decomposition/C/N not measured | WOS |
| 379 | Neves et al. | 2010 | Canopy Herbivory and Insect Herbivore Diversity in a Dry Forest-Savanna Transition in Brazil | Biotropica | 42(1): 112-118 | Decomposition/C/N not measured | WOS |
| 380 | Oxbrough et al. | 2010 | Ground-dwelling invertebrates in reforested conifer plantations | Forest Ecology and Management | 259(10): 2111-2121 | No exclusion of invertbrates | WOS |
| 381 | Wegener and Alberti | 2010 | Effects of a windthrow event in the forest of the peninsula Darss on the gamasid fauna (Arachnida) and Collembola | Trends of Acarology (Book) | 117-121 | Decomposition/C/N not measured | WOS |
| 382 | Oelbermann and Scheu | 2010 | Trophic guilds of generalist feeders in soil animal communities as indicated by stable isotope analysis (N-15/N-14) | Bulletin of Entomological Research | 100(5): 511-20 | Decomposition/C/N not measured | WOS |
| 383 | Godbold et al. | 2009 | Consumer and resource diversity effects on marine macroalgal decomposition | Oikos | 118(1): 77 - 86 | Aquatic system | WOS |
| 384 | Taylor | 2009 | Biological Flora of the British Isles: Urtica dioica L. | Journal of Ecology | 97(6): 1436-1458 | Review/meta-analysis |  |
| 385 | Knapp et al. | 2009 | Diet-related composition of the gut microbiota of Lumbricus rubellus as revealed by a molecular fingerprinting technique and cloning | Soil Biology and Biochemistry | 41(11): 2299-2307 | Diet study of microbiota | WOS |
| 386 | Cooney and Simon | 2009 | Influence of dissolved organic matter and invertebrates on the function of microbial films in groundwater | Microbial Ecology | 58(4): 968 | Aquatic system | WOS |
| 387 | Younginger et al. | 2009 | Interactive effects of mycorrhizal fungi, salt stress, and competition on the herbivores of Baccharis halimifolia | Ecological Entomology | 34(5): 580-587 | Decomposition/C/N not measured | WOS |
| 388 | Ayres et al. | 2009 | Tree Species Traits Influence Soil Physical, Chemical, and Biological Properties in High Elevation Forests | PLoS ONE | 4(6): e5964 | No exclusion of invertbrates | WOS |
| 389 | Brys and Jacquemyn | 2009 | Biological Flora of the British Isles: Primula veris L. | Journal of Ecology | 97(3): 581-600 | Review/meta-analysis | WOS |
| 390 | Bonkowski et al. | 2009 | Rhizosphere fauna: the functional and structural diversity of intimate interactions of soil fauna with plant roots | Plant and Soil | 321(1): 213-233 | Review/meta-analysis | WOS |
| 391 | Farji-Brener et al. | 2009 | Small-scale disturbances spread along trophic chains: leaf-cutting ant nests, plants, aphids, and tending ants | Ecological Research | 24(1): 139–145 | Decomposition/C/N not measured | WOS |
| 392 | Borowicz | 2009 | Organic Farm Soil Improves Strawberry Growth But Does Not Diminish Spittlebug Damage | Journal of Sustainable Agriculture | 33(2): 177-188 | No exclusion of invertbrates | WOS |
| 393 | Telford et al. | 2009 | Bioaccumulation of antimony and arsenic in a highly contaminated stream adjacent to the Hillgrove Mine, NSW, Australia | Environmental Chemistry | 6(2): 133-143 | Aquatic system | WOS |
| 394 | Towns et al. | 2009 | Predation of seabirds by invasive rats: multiple indirect consequences for invertebrate communities | Oikos | 118(3): 420-430 | No exclusion of invertbrates | WOS |
| 395 | Doblas-Miranda et al. | 2009 | Different microhabitats affect soil macroinvertebrate assemblages in a Mediterranean arid ecosystem | Applied Soil Ecology | 41(3): 329-335 | No exclusion of invertbrates | WOS |
| 396 | Knapp et al. | 2009 | Molecular fingerprinting analysis of the gut of Cylindroiulus fulviceps (Diplopoda) | Pedobiologia | 52(5): 325-336 | Methods paper | WOS |
| 397 | Salamon and Zaitsev | 2008 | Soil macrofaunal response to forest conversion from pure coniferous stands into semi-natural montane forests | Applied Soil Ecology | 40(3): 491-498 | No exclusion of invertbrates | WOS |
| 398 | Dunham | 2008 | Above and belowground impacts of terrestrial mammals and birds in a tropical forest | Oikos | 117(4): 571-579 | Decomposition/C/N not measured | WOS |
| 399 | Schuldt et al. | 2008 | Communities of ground-living spiders in deciduous forests: does tree species diversity matter? | Biodiversity and Conservation | 17(5): 1267-1284 | Decomposition/C/N not measured | WOS |
| 400 | Icoz and Stotzky | 2008 | Fate and effects of insect-resistant Bt crops in soil ecosystems | Soil Biology and Biochemistry | 40(3): 559-586 | Review/meta-analysis | WOS |
| 401 | Schweitzer et al. | 2008 | From genes to ecosystems: the genetic basis of condensed tannins and their role in nutrient regulation in a populus model system | Ecosystems | 11(6): 1005-1020 | Review/meta-analysis | WOS |
| 402 | Cannicci et al. | 2008 | Faunal impact on vegetation structure and ecosystem function in mangrove forests: A review | Aquatic Botany | 89(2): 186-200 | Review/meta-analysis | WOS |
| 403 | Kaplan et al. | 2008 | Constitutive and induced defenses to herbivory in above- and belowground plant tissues | Ecology | 89(2): 392-406 | Review/meta-analysis | WOS |
| 404 | Risch et al. | 2008 | Abundance and distribution of organic mound-building ants of the Formica rufa group in Yellowstone National Park | Journal of Applied Entomology | 132(4): 326-336 | Decomposition/C/N not measured | WOS |
| 405 | Jurgensen et al. | 2008 | Organic mound-building ants: their impact on soil properties in temperate and boreal forests | Journal of Applied Entomology | 132(4): 266-275 | Review/meta-analysis | WOS |
| 406 | Halaj et al. | 2008 | Responses of litter-dwelling spiders and carabid beetles to varying levels and patterns of green-tree retention | Forest Ecology and Management | 255(3): 887-900 | Decomposition/C/N not measured | WOS |
| 407 | Schon et al. | 2008 | Soil fauna in grazed New Zealand hill country pastures at two management intensities | Applied Soil Ecology | 40(2): 218-228 | Management impacts on soil invertebrates | WOS |
| 408 | Leger et al. | 2007 | The interaction between soil nutrients and leaf loss during early 14 establishment in plant invasion | Forest Science | 53(6): 701-709 | Decompositon/C/N not measured | WOS |
| 409 | Manning et al. | 2007 | Saprotrophy of Conidiobolus and Basidiobolus in leaf litter | Mycological Research | 111(12): 1437-1449 | No invertebrates studied | WOS |
| 410 | Paoletti et al. | 2007 | Detritivores as indicators of landscape stress and soil degradation | Australian Journal of Experimental Agriculture | 47(4): 412-423 | Decomposition/C/N not measured | WOS |
| 411 | Majer et al. | 2007 | Invertebrates and the restoration of a forest ecosystem: 30 years of research following bauxite mining in Western Australia | Restoration Ecology | 15(s4): S104-S115 | No exclusion of invertbrates | WOS |
| 412 | McGlynn et al. | 2007 | Phosphorus limits tropical rain forest litter fauna | Biotropica | 39(1): 50-53 | No exclusion of invertbrates | WOS |
| 413 | Meehan and Lindroth | 2007 | Modeling nitrogen flux by larval insect herbivores from a temperate hardwood forest | Oecologia | 153(4): 833-843 | Modeling paper | WOS |
| 414 | Classen et al. | 2007 | Season mediates herbivore effects on litter and soil microbial abundance and activity in a semi-arid woodland | Plant and Soil | 295(1): 217-227 | Decomposition/C/N not measured | WOS |
| 415 | Curry and Olaf | 2007 | The feeding ecology of earthworms - A review | Pedobiologia | 50(6): 463-477 | Review/meta-analysis | WOS |
| 416 | Dalrymple | 2007 | Biological flora of the British Isles: Melampyrum sylvaticum L. | Journal of Ecology | 95(3): 583-597 | Review/meta-analysis | WOS |
| 417 | Huang et al. | 2007 | Toads (Bufo bankorensis) influence litter chemistry but not litter invertebrates and litter decomposition rates in a subtropical forest of Taiwan | Journal of Tropical Ecology | 23(2): 161-168 | Toad exclusion; not invertebrates | WOS |
| 418 | Yeates | 2007 | Abundance, diversity, and resilience of nematode assemblages in forest soils | Canadian Journal of Forest Research | 37(2): 216-225 | Decomposition/C/N not measured | WOS |
| 419 | Krivtsov et al. | 2006 | Ecological study of the forest litter meiofauna of a unique Scottish woodland | Animal Biology | 56(1): 69-93 | Decomposition/C/N not measured | WOS |
| 420 | Meehan et al. | 2006 | Energetic equivalence in a soil arthropod community from an aspen-conifer forest | Pedobiologia | 50(4): 307-312 | Decomposition/C/N not measured | WOS |
| 421 | Salamon et al. | 2006 | Transitory dynamic effects in the soil invertebrate community in a temperate deciduous forest: Effects of resource quality | Soil Biology and Biochemistry | 38(2): 209-221 | No exclusion of invertbrates | WOS |
| 422 | Albers et al. | 2006 | Incorporation of plant carbon into the soil animal food web of an arable system | Ecology | 87(1): 235-245 | Decomposition/C/N not measured | WOS |
| 423 | Woodcock et al. | 2006 | Land-use effects on catchment- and patch-scale habitat and macroinvertebrate responses in the Adirondack uplands | American Fisheries Society Symposium | 48: 395–411 | Aquatic system | WOS |
| 424 | Ammer et al. | 2006 | Factors influencing the distribution and abundance of earthworm communities in pure and converted Scots pine stands | Applied Soil Ecology | 33(1): 10-21 | Decomposition/C/N not measured | WOS |
| 425 | Rombke et al. | 2006 | Identification of potential organisms of relevance to Canadian boreal forest and northern lands for testing of contaminated soils | Environmental Reviews | 14(2): 137-167 | Review/meta-analysis | WOS |
| 426 | Fine et al. | 2006 | The growth-defense trade-off and habitat specialization by plants in Amazonian forests | Ecology | 87(sp7): S150-S162 | Invertebrates not studied | WOS |
| 427 | Pritekel et al. | 2006 | Impacts from invasive plant species and their control on the plant community and belowground ecosystem at Rocky Mountain National Park, USA | Applied Soil Ecology | 32(1): 132-141 | Studied impacts of an invasive plant | WOS |
| 428 | Stadler et al. | 2006 | The ecology of energy and nutrient fluxes in hemlock forests invaded by hemlock woolly adelgid | Ecology | 87(7): 1792-1804 | Studied impacts of an invasive plant | WOS |
| 429 | Blouin et al. | 2005 | Belowground organism activities affect plant aboveground phenotype, inducing plant tolerance to parasites | Ecology Letters | 8(2): 202-208 | Decomposition/C/N not measured | WOS |
| 430 | Walton and Steckler | 2005 | Contrasting effects of salamanders on forest-floor macro- and mesofauna in laboratory microcosms | Pedobiologia | 49(1): 51-60 | Salamander effects; Decomposition/C/N was not measured | WOS |
| 431 | Migge-Kleian et al. | 2005 | Impact of forest distrubance and land use change on soil and litter arthropod assemblages in tropical rainforest margins | Stability of Tropical Rainforest Margins (book) | 147-163 | Decomposition/C/N not measured | WOS |
| 432 | McNeil and Cushman | 2005 | Indirect effects of deer herbivory on local nitrogen availability in a coastal dune ecosystem | Oikos | 110(1): 124-132 | No invertebrates studied | WOS |
| 433 | Cook and Dawes-Gromadzki | 2005 | Stable isotope signatures and landscape functioning in banded vegetation in arid-central Australia | Landscape Ecology | 20(6): 649–660 | No exclusion of invertbrates | WOS |
| 434 | Schneider and Maraun | 2005 | Feeding preferences among dark pigmented fungal taxa ("Dematiacea") indicate limited trophic niche differentiation of oribatid mites (Oribatida, Acari) | Pedobiologia | 49(1): 61-67 | Decomposition/C/N not measured | WOS |
| 435 | Krivtsov et al. | 2005 | Forest litter bacteria: relationships with fungi, microfauna, and litter composition over a winter-spring period | Polish Journal of Ecology | 53(3): 383–394 | No exclusion of invertbrates |  |
| 436 | Mohr et al. | 2005 | Wild boar and red deer affect soil nutrients and soil biota in steep oak stands of the Eifel | Soil Biology and Biochemistry | 47(4): 693-700 | No exclusion of invertbrates | WOS |
| 437 | Halaj et al. | 2005 | Trophic structure of a macroarthropod litter food web in managed coniferous forest stands: a stable isotope analysis with delta N-15 and delta C-13 | Pedobiologia | 49(2): 109-118 | Decomposition/C/N not measured | WOS |
| 438 | Fonseca et al. | 2005 | Flower-heads, herbivores, and their parasitoids: food web structure along a fertility gradient | Ecological Entomology | 30(1): 36-46 | Decomposition/C/N not measured | WOS |
| 439 | Fagan et al. | 2004 | Spatially structured herbivory and primary succession at Mount St Helens: field surveys and experimental growth studies suggest a role for nutrients | Ecological Entomology | 29(4): 398-409 | Decomposition/C/N not measured | WOS |
| 440 | Hamilton et al. | 2004 | Insect herbivory in an intact forest understory under experimental CO2 enrichment | Oecologia | 138(4): 566-573 | Decomposition/C/N not measured | WOS |
| 441 | Stockan et al. | 2004 | Phytochemistry of Scots pine as a driver of spatial heterogeneity in insect herbivore and broader diversity in a native woodland landscape | Proceedings of the twelfth annual IALE (UK) conference | 340-343 | No exclusion of invertbrates | WOS |
| 442 | Veldtman and McGeoch | 2003 | Gall-forming insect species richness along a non-scleromorphic vegetation rainfall gradient in South Africa: The importance of plant community composition | Austral Ecology | 28(1): 1-13 | No exclusion of invertbrates | WOS |
| 443 | Mitchell | 2003 | Trophic control of grassland production and biomass by pathogens | Ecology Letters | 6(2): 147-155 | No invertebrates studied | WOS |
| 444 | Sadaka and Ponge | 2003 | Climatic effects on soil trophic networks and the resulting humus profiles in holm oak (Quercus rotundifolia) forests in the High Atlas of Morocco as revealed by correspondence analysis | Soil Science | 54(4): 767-777 | Decomposition/C/N not measured | WOS |
| 445 | Bardgett and Wardle | 2003 | Herbivore-mediated linkages between aboveground and belowground communities | Ecology | 84(9): 2258-2268 | Review/meta-analysis | WOS |
| 446 | Reimchen et al. | 2003 | Isotopic evidence for enrichment of salmon-derived nutrients in vegetation, soil, and insects in Riparian zones in coastal British Columbia | American Fisheries Society Symposium | 34:59-70 | Impacts of salmon on soil invertebrates; decomposition/C/N not measured | WOS |
| 447 | Pokarzhevskii et al. | 2003 | Microbial links and element flows in nested detrital food webs | Pedobiologia | 47(3): 213-224 | Review/meta-analysis | WOS |
| 448 | Fellerhoff et al. | 2003 | Stable carbon and nitrogen isotope signatures of decomposition tropical macrophytes | Aquatic Ecology | 37(4): 361–375 | Aquatic system | WOS |
| 449 | Beard et al. | 2003 | The effects of the frog Eleutherodactylus coqui on invertebrates and ecosystem processes at two scales in the Luquillo Experimental Forest, Puerto Rice | Journal of Tropical Ecology | 19(6): 607-617 | Studied frog impacts on Decomposition/C/N | WOS |
| 450 | Scheu et al. | 2003 | The soil fauna community in pure and mixed stands of beech and spruce of different age: trophic structure and structuring forces | Oikos | 101(2): 225-238 | Decomposition/C/N not measured | WOS |
| 451 | Liiri et al. | 2002 | Community composition of soil microarthropods of acid forest soils as affected by wood ash application | Pedobiologia | 46(2): 108-124 | Only diversity of invertebrates measured; Decomposition/C/N was not measured | WOS |
| 452 | Ferguson and Do | 2002 | Dynamics of springtail and mite populations: the role of density dependence, predation, and weather | Ecological Entomology | 27(5): 565-573 | Only population dynamics studied; Decomposition/C/N was not measured | WOS |
| 453 | Lavelle et al. | 2002 | Functional domains in soils | Ecological Research | 17(4): 441-450 | Review/meta-analysis | WOS |
| 454 | Stiling et al. | 2002 | Elevated atmospheric CO2 lowers herbivore abundance, but increases leaf abscission rates | Global Change Biology | 8(7): 658-667 | Decomposition/C/N not measured | WOS |
| 455 | Goverde et al. | 2002 | Species-specific reactions to elevated CO2 and nutrient availability in four grass species | Basic and Applied Ecology | 3(3): 221-227 | No invertebrate exclusion | WOS |
| 456 | Moore et al. | 2002 | Effects of two silvicultural practices on soil fauna abundance in a northern hardwood forest, Quebec, Canada | Canadian Journal of Soil Science | 82(1): 105-113 | No invertebrate exclsusion | WOS |
| 457 | Richardson et al. | 2002 | How do nutrients and warming impact on plant communities and their insect herbivores? A 9-year study from a sub-Arctic heath | Journal of Ecology | 90(3): 544-556 | Decomposition/C/N not measured | WOS |
| 458 | Frouz et al. | 2002 | The potential effect of high atmospheric CO2 on soil fungi-invertebrate interactions | Global Change Biology | 8(4): 339-344 | Decomposition/C/N not measured | WOS |
| 459 | Pickett et al. | 2001 | Urban ecological systems: Linking terrestrial ecological, physical, and socioeconomic components of metropolitan areas | Annual Review of Ecology and Systematics | 32(1): 127-157 | Review/meta-analysis | WOS |
| 460 | Peterson et al. | 2001 | Single-shrub influence on earthworms and soil macroarthropods in the southern California chaparral | Pedobiologia | 45(6): 509-522 | Decomposition/C/N not measured | WOS |
| 461 | Belnap and Phillips | 2001 | Soil biota in an ungrazed grassland: Response to annual grass (Bromus tectorum) invasion | Ecological Applications | 11(5): 261-1275 | Decomposition/C/N not measured | WOS |
| 462 | Zaitsev and van Straalen | 2001 | Species diversity and metal accumulation in oribatid mites (Acari, Oribatida) of forests affected by a metallurgical plant | Pedobiologia | 45(5): 467-479 | Decomposition/C/N not measured | WOS |
| 463 | Niwa | 2001 | Soil, litter, and coarse woody debris habitats for arthropods in eastern Oregon and Washington | Northwest Science | 75: 141-148 | Decomposition/C/N not measured | WOS |
| 464 | Osler et al. | 2000 | Changes in free living soil nematode and microarthropod communities under a conola-wheat-lupin rotation in Western Australia | Soil Research | 38(1): 47-60 | Decomposition/C/N not measured | WOS |
| 465 | Buckland and Grime | 2000 | The effects of trophic structure and soil fertility on the assembly of plant communities: a microcosm experiment | Oikos | 91(2): 336-352 | Decomposition/C/N not measured | WOS |
| 466 | Lavelle | 2000 | Ecological challenges for soil science | Soil Science | 165(1): 73-86 | Review/meta-analysis | WOS |
| 467 | Blum et al. | 2000 | Changes in Sr/Ca, Ba/Ca and Sr-87/Sr-86 ratios between trophic levels in two forest ecosystems in the northeastern USA | Biogeochemistry | 49(1): 87-101 | Decomposition/C/N not measured | WOS |
| 468 | Bis et al. | 2000 | Effects of catchment properties on hydrochemistry, habitat complexity and invertebrate community structure in a lowland river | Assessing the Ecological Integrity of Running Waters (Book) | 369-387 | Aquatic system | WOS |
| 469 | Johnson | 2000 | The contribution of microarthropods to aboveground food webs: A review and model of belowground transfer in a coniferous forest | The American Midland Naturalist | 143(1): 226-238 | Review/meta-analysis | WOS |
| 470 | Van Auken | 2000 | Shrub invasions of North American semiarid grasslands | Annual Review of Ecology and Systematics | 31(1): 197-215 | Review/meta-analysis | WOS |
| 471 | Horne | 2000 | Phytoremediation by constructed wetlands | Phytoremediation of Contaminated Soil and Water (Book) | 1: 13-39 | No exclusion of invertbrates | WOS |
| 472 | Wolters | 2000 | Invertebrate control of soil organic matter stability | Biology and Fertility of Soils | 31(1): 1-19 | Review/meta-analysis | WOS |
| 473 | Anaya | 1999 | Allelopathy as a tool in the management of biotic resources in agroecosystems | Critical Reviews in Plant Sciences | 18(6): 697-739 | Decomposition/C/N not measured | WOS |
| 474 | Wardle et al. | 1999 | Effects of agricultural intensification on soil-associated arthropod population dynamics, community structure, diversity, and temporal variability over a seven-year period | Soil Biology and Biochemistry | 31(12): 1691-1706 | Decomposition/C/N not measured | WOS |
| 475 | Zheng et al. | 1999 | How do soil organisms affect total organic nitrogen storage and substrate nitrogen to carbon ratio in soils? A theoretical analysis | Oikos | 86(3): 430-442 | Theoretical paper | WOS |
| 476 | Koehler | 1999 | Predatory mites (Gamasina, Mesostigmata) | Agriculture, Ecosystems & Environment | 74(1-3): 395-410 | Review/meta-analysis | WOS |
| 477 | Behan-Pelletier | 1999 | Oribatid mite biodiversity in agroecosystems: role for bioindication | Agriculture, Ecosystems & Environment | 74(1-3): 411-423 | Review/meta-analysis | WOS |
| 478 | Fraser and Grime | 1999 | Interacting effects of herbivory and fertility on a synthesized plant community | Journal of Ecology | 87(3): 514-525 | Decomposition/C/N not measured | WOS |
| 479 | Frouz | 1999 | Use of soil dwelling Diptera (Insecta, Diptera as bioindicators: a review of ecological requirements and response to disturbance | Agriculture, Ecosystems & Environment | 74(1-3): 167-186 | Review/meta-analysis | WOS |
| 480 | Chambers et al. | 1999 | Seed and seedling ecology of pinon and juniper species in the pygmy woodlands of western North America | The Botanical Review | 65(1): 1-38 | Review/meta-analysis | WOS |
| 481 | James et al. | 1999 | Provision of watering points in the Australian arid zone: a review of effects on biota | Journal of Arid Environments | 41(1): 87-121 | Review/meta-analysis | WOS |
| 482 | Kovaliukh et al. | 1998 | C-14 cycle in the hot zone around Chernobyl | Radiocarbon | 40(1): 391-397 | Radiative impacts on carbon dynamics; Decomposition/C/N was not measured | WOS |
| 483 | Gers | 1998 | Diversity of energy fluxes and interactions between arthropod communities: from soil to cave | Acta Oecologia | 19(3): 205-213 | Decomposition/C/N not measured | WOS |
| 484 | Laarkso and Setala | 1998 | Composition and trophic structure of detrital food web in ant nest mounds of Formica aquilonia and in the surrounding forest soil | Oikos | 81(2): 266-278 | Decomposition/C/N not measured | WOS |
| 485 | Erelli et al. | 1998 | Altitudinal patterns in host suitability for forest insects | Oecologia | 117(1-2): 133-142 | No exlusion of invertebrates | WOS |
| 486 | Giller et al. | 1997 | Agricultural intensification, soil biodiversity, and agroecosystem function | Applied Soil Ecology | 6(1): 3-16 | Review/meta-analysis | WOS |
| 487 | Perry et al. | 1997 | Response of soil and leaf litter microarthropods to forest application of diflubenzuron | Ecotoxicology | 6(2): 87-99 | Studied the effects of diflubenzuron on soil and litter arthropods | WOS |
| 488 | Paquin and Coderre | 1997 | Decomposition of genetically engineered tobacco under field conditions: Persistence of the proteinase inhibitor I product and effects on soil microbial respiration and protozoa, nematode and microarthropod populations | Journal of Applied Ecology | 34(3): 767-777 | Decomposition/C/N not measured | WOS |
| 489 | Orians and Floyd | 1997 | The susceptibility of parental and hybrid willows to plant enemies under contrasting soil nutrient conditions | Oecologia | 109(3): 407-413 | Decomposition/C/N not measured | WOS |
| 490 | Ardon | 1997 | The impact of peat cutting on fauna and flora | Landscape Archaeology an Ecology | 4: 10-27 | Review/meta-analysis | WOS |
| 491 | Lavelle et al. | 1997 | Soil function in a changing world: the role of invertebrate ecosystem engineers | European Journal of Soil Biology | 33(4): 159-193 | Review/meta-analysis | WOS |
| 492 | Cook et al. | 1996 | Safety of microorganisms intended for pest and plant disease control: A framework for scientific evaluation | Biological Control | 7(3): 333-351 | Review/meta-analysis | WOS |
| 493 | Sleaford et al. | 1996 | A pilot analysis of gut contents in termites from the Mbalmayo forest reserve, Cameroon | Ecological Entomology | 21(3): 279-288 | A gut analysis of termites | WOS |
| 494 | Dejean et al. | 1996 | Ants inhabiting Cubitermes termitaries in African rain forests | Biotropica | 28(4):701-713 | Decomposition/C/N not measured | WOS |
| 495 | Edwards | 1996 | New insights into early land ecosystems: a glimpse of a Lilliputian world | Review of Palaeobotany and Palynology | 90(3-4): 159-174 | Review/meta-analysis | WOS |
| 496 | Stone et al. | 1996 | Canopy microfungi: function and diversity | Northwest Science | 70: 37-45 | A characterization of canopy microfungi | WOS |
| 497 | Poinsotbalaguer and Tabone | 1995 | Impact of chronic gamma-irradiation on the litter decay of a mixed mediterranean forest in cadarache - France microarthropods response | Pedobiologia | 39(4): 344-350 | A gamma-radiation experiment; no animal exclusion | WOS |
| 498 | Hoekstra et al. | 1995 | Soil arthropod abundance in coast redwood forest - effect of selective timber harvest | Environmental Entomology | 24(2): 246-252 | Only measured invertebrate abundances; no Decomposition/C/N or exclusion measured | WOS |
| 499 | Callaghan and Jonasson | 1995 | Arctic terrestrial ecosystems and environmental change | Philosophical Transactions of the Royal Society of London | 352(1699): 259-276 | Review/meta-analysis | WOS |
| 500 | Yeates | 1995 | Effect of sewage effluent on soil fauna in a pinus-radiata plantation | Soil Research | 33(3): 555-564 | Decomposition/C/N not measured | WOS |
| 501 | Blair et al. | 1994 | A comparison of the forest floor invertebrate communities of 4 forest types in the northeastern United States | Pedobiologia | 38(2): 146-160 | No invertebrate exclusion or evaluation of Decomposition/C/N | WOS |
| 502 | Langenheim | 1994 | Higher plant terpenoids-A phytocentric overview of their ecological roles | Journal of Chemical Ecology | 20(6): 1223-1280 | Review/meta-analysis | WOS |
| 503 | Pouyat et al. | 1994 | Environmental effects of forest soil invertebrate and fungal densities in oak stands along an urban-rural land-use gradient | Pedobiologia | 38(5): 385-399 | No exclusion of arthropods/invertebrates; only densities measured | WOS |
| 504 | Roth | 1993 | Investigations on lead in the soil invertebrates of a forest ecosystem | Pedobiologia | 37(5): 270-279 | No exclusion of arthropods/invertebrates; only densities measured | WOS |
| 505 | Parmelee et al. | 1993 | Soil microcosm for testing the effects of chemical pollutants on soil fauna communities and trophic structure | Environmental Toxicology and Chemistry | 12(8): 1477-1486 | No Decomposition/C/N measured and no invertebrate exclusion | WOS |
| 506 | Lattin | 1993 | Arthropod diversity and conservation in old-growth Northwest forests | American Zoologist | 33(6): 578-587 | Review/meta-analysis | WOS |
| 507 | Sweeney | 1993 | Effects of streamside vegetation on macroinvertebrate communities of White Clay Creek in Eastern North America | Proceedings of the Academy of Natural Sciences of Philadelphia | 144: 291-340 | Aquatic system | WOS |
| 508 | Stork and Blackburn | 1993 | Abundance, body-size, and biomass of arthropods in tropical forest | Oikos | 67(3): 483-489 | Decomposition/C/N not measured | WOS |
| 509 | Kikkawa and Dwey | 1992 | Use of scattered resources in rain forest of humid tropic lowlands | Biotropica | 24(2B): 293-308 | No Decomposition/C/N measured and no invertebrate exclusion | WOS |
| 510 | Facelli and Pickett | 1991 | Plant litter: It Dynamics and Effects on Plant Community Structure | The Botanical Review | 57(1): 1-32 | Review/meta-analysis | WOS |
| 511 | Robertson | 1991 | Plant-animal interactions and the structure and function of mangrove forest ecosystems | Australian Journal of Ecology | 16(4): 433-443 | No exlusion of invertebrates | WOS |
| 512 | Wolters | 1991 | Soil invertebrates: Effects on nutrient turnover and soil structure-A review | Zeitschrift für Pflanzenernährung und Bodenkunde | 154(6): 389-402 | Review/meta-analysis | WOS |
| 513 | Cocking et al. | 1991 | Compartmentalization of mercury in biotic components of terrestrial flood-plain ecosystems adjacent to the South River at Waynesboro, VA | Water, Air, and Soil Pollution | 57(1): 159-170 | No exclusion of invertbrates | WOS |

**Table S2.** The screening of articles from the second round after full article evaluation. WOS = Web of Science.

| **Study ID** | **Author** | **Year** | **Title** | **Journal** | **Reason for exclusion** | **Source** |
| --- | --- | --- | --- | --- | --- | --- |
| 267 | Bisanzio et al. | 2021 | Arboviral diseases and poverty in Alabama, 2007-2017 | Plos Neglected Tropical Disease | No data on arboviral disease in low vs. high SES | WOS |
| 268 | Uelmen et al. | 2021 | Dynamics of data availability in disease modeling: An example evaluating the trade-offs of ultra-fine-scale factors applied to human West Nile virus disease models in the Chicago area, USA | Plos One | No mosquito abundance data in low vs. high SES | WOS |
| 269 | Rothman et al. | 2021 | Higher West Nile Virus Infection in Aedes albopictus (Diptera: Culicidae) and Culex (Diptera: Culicidae) Mosquitoes From Lower Income Neighborhoods in Urban Baltimore, MD | Journal of Medical Entomology | Dataset already represented | WOS |
| 270 | Humphreys et al. | 2021 | Vector Surveillance, Host Species Richness, and Demographic Factors as West Nile Disease Risk Indicators | Viruses | No mosquito abundance data in low vs. high SES | WOS |
| 271 | Yee et al. | 2021 | Mosquitoes (Diptera: Culicidae) on the islands of Puerto Rico and Vieques, U.S.A. | Acta Tropica | No mosquito abundance data in low vs. high SES | Supplemental |
| 272 | Watts et al. | 2020 | Infuence of socio-economic, demographic and climate factors on the regional distribution of dengue in the United States and Mexico | International Journal of Health Geographics | No mosquito abundance data in low vs. high SES | WOS |
| 273 | Butterworth | 2020 | ‘Clean up your rain gutters!’: mosquito control, responsibility, and blame following the 2009–2010 dengue fever outbreak in Key West, Florida | GeoJournal | No mosquito abundance data in low vs. high SES | WOS |
| 274 | Hinojosa et al. | 2020 | Detection of a Locally-Acquired Zika Virus Outbreak in Hidalgo County, Texas through Increased Antenatal Testing in a High-Risk Area | Tropical Medicine and Infectious Diseases | No mosquito abundance data in low vs. high SES | WOS |
| 275 | Rohat et al. | 2020 | Intersecting vulnerabilities: climatic and demographic contributions to future population exposure toAedes-borne viruses in the United States | Environmental Research Letters | Future projections; not based on past collections | WOS |
| 276 | Kala et al. | 2020 | Exploring the socio-economic and environmental components of infectious diseases using multivariate geovisualization: West Nile Virus | PeerJ | No mosquito abundance data in low vs. high SES | WOS |
| 277 | Lemanski et al. | 2020 | Coordination among neighbors improves the efficacy of Zika control despite economic costs | Plos Neglected Tropical Disease | No mosquito abundance data in low vs. high SES | WOS |
| 278 | Graves et al. | 2020 | Demographic, socioeconomic and disease knowledge factors, but not population mobility, associated with lymphatic filariasis infection in adult workers in American Samoa in 2014 | Parasites and Vectors | No mosquito abundance data in low vs. high SES | WOS |
| 279 | Katz et al. | 2020 | Aedes albopictus Body Size Differs Across Neighborhoods With Varying Infrastructural Abandonment | Journal of Medical Entomology | No mosquito abundance data in low vs. high SES | WOS |
| 280 | Beaulieu et al. | 2020 | Mosquito diversity and dog heartworm prevalence in suburban areas | Parasites and Vectors | No mosquito abundance data in low vs. high SES | WOS |
| 281 | Myer et al. | 2020 | Mapping Aedes aegypti (Diptera: Culicidae) and Aedes albopictus Vector Mosquito Distribution in Brownsville, TX | Journal of Medical Entomology | No mosquito abundance data in low vs. high SES | WOS |
| 282 | Borucki et al. | 2020 | Multiscale analysis for patterns of Zika virus genotype emergence, spread, and consequence | Plos One | No mosquito abundance data in low vs. high SES | WOS |
| 283 | Crespo et al. | 2019 | Linking Wetland Ecosystem Services to Vector-borne Disease: Dengue Fever in the San Juan Bay Estuary, Puerto Rico | Wetlands | No mosquito abundance data in low vs. high SES | WOS |
| 284 | Wilke et al. | 2019 | Community Composition and Year-round Abundance of Vector Species of Mosquitoes make Miami-Dade County, Florida a Receptive Gateway for Arbovirus entry to the United States | Scientific Reports | No mosquito abundance data in low vs. high SES | WOS |
| 285 | Chen et al. | 2019 | An operational machine learning approach to predict mosquito abundance based on socioeconomic and landscape patterns | Landscape Ecology | Future projections; not based on past collections | WOS |
| 286 | Bodner et al. | 2019 | Relationships Among Immature-Stage Metrics and Adult Abundances of Mosquito Populations in Baltimore, MD | Journal of Medical Entomology | Dataset already represented | WOS |
| 287 | Whiteman et al. | 2019 | A Novel Sampling Method to Measure Socioeconomic Drivers of Aedes albopictus Distribution in Mecklenburg County, North Carolina | Environmental Research and Public Health | No mosquito abundance data in low vs. high SES | WOS |
| 288 | Yee et al. | 2019 | Linking Water Quality to Aedes aegypti and Zika in Flood-Prone Neighborhoods | Ecohealth | No mosquito abundance data in low vs. high SES | Supplemental |
| 289 | Romeo-Aznar et al. | 2018 | Mosquito-borne transmission in urban landscapes: the missing link between vector abundance and human density | Proceedings of the Royal Society B. | Data not from the US; no mosquito abundance data in low vs. high SES | WOS |
| 290 | Monaghan et al. | 2018 | The potential impacts of 21st century climatic and population changes on human exposure to the virus vector mosquito Aedes aegypti | Climatic Change | Future projections; not based on past collections | WOS |
| 291 | Gardner et al. | 2018 | Inferring the risk factors behind the geographical spread and transmission of Zika in the Americas | Plos One Neglected Tropical Diseases | No mosquito abundance data in low vs. high SES | WOS |
| 292 | Piltch-Loeb et al. | 2017 | Risk salience of a novel virus: US population risk perception, knowledge, and receptivity to public health interventions regarding the Zika virus prior to local transmission | Plos One | No mosquito abundance data in low vs. high SES | WOS |
| 293 | Little et al | 2017 | Local environmental and meteorological conditions influencing the invasive mosquito Ae. albopictus and arbovirus transmission risk in New York City | Plos One Neglected Tropical Diseases | No mosquito abundance data in low vs. high SES | WOS |
| 294 | Sallam et al. | 2017 | Spatio-Temporal Distribution of Vector-Host Contact (VHC) Ratios and Ecological Niche Modeling of the West Nile Virus Mosquito Vector, Culex quinquefasciatus, in the City of New Orleans, LA, USA | Environmental Research and Public Health | No mosquito abundance data in low vs. high SES | WOS |
| 295 | Zhang et al. | 2017 | Spread of Zika virus in the Americas | Proceedings of the National Academy of Sciences | Modeling; study conducted outside the US | WOS |
| 296 | Mordecai et al. | 2017 | Detecting the impact of temperature on transmission of Zika, dengue, and chikungunya using mechanistic models | Plos Neglected Tropical Disease | Modeling paper | WOS |
| 297 | Lockaby et al. | 2016 | Climatic, ecological, and socioeconomic factors associated with West Nile virus incidence in Atlanta, Georgia, USA | Journal of Vector Ecology | No mosquito abundance data in low vs. high SES | WOS |
| 298 | Skaff and Cheruvelil | 2016 | Fine-scale wetland features mediate vector and climatedependent macroscale patterns in human West Nile virus incidence | Landscape Ecology | No mosquito abundance data in low vs. high SES | WOS |
| 299 | Santos-Vega et al. | 2016 | Climate forcing and infectious disease transmission in urban landscapes: integrating demographic and socioeconomic heterogeneity | Annals of New York Academy of Sciences | No mosquito abundance data in low vs. high SES | WOS |
| 300 | Vitek et al. | 2014 | Dengue Vectors, Human Activity, and Dengue Virus Transmission Potential in the Lower Rio Grande Valley, Texas, United States | Journal of Medical Entomology | No mosquito abundance data in low vs. high SES | WOS |
| 301 | Blake and Garcia-Blanco | 2014 | Human Genetic Variation and Yellow Fever Mortality during 19th Century US Epidemics | MBIO | No mosquito abundance data in low vs. high SES | WOS |
| 302 | Faraji et al. | 2014 | Comparative Host Feeding Patterns of the Asian Tiger Mosquito, Aedes albopictus, in Urban and Suburban Northeastern USA and Implications for Disease Transmission | Plos Neglected Tropical Disease | No mosquito abundance data in low vs. high SES | Supplemental |
| 303 | Fonseca et al. | 2013 | Area-wide management of Aedes albopictus. Part 2: Gauging the efficacy of traditional integrated pest control measures against urban container mosquitoes | Pest Management Science | Dataset already represented | WOS |
| 304 | Dowling et al. | 2013 | Socioeconomic Status Affects Mosquito (Diptera: Culicidae) Larval Habitat Type Availability and Infestation Level | Journal of Medical Entomology | No mosquito abundance data in low vs. high SES | WOS |
| 305 | Eisen and Moore | 2013 | Aedes (Stegomyia) aegypti in the Continental United States: A Vector at the Cool Margin of Its Geographic Range | Journal of Medical Entomology | Review | WOS |
| 306 | Dowling et al. | 2013 | Linking Mosquito Infestation to Resident Socioeconomic Status, Knowledge, and Source Reduction Practices in Suburban Washington, DC | Ecohealth | No mosquito abundance data in low vs. high SES | WOS |
| 307 | Reiter et al. | 2013 | Texas lifestyle limits transmission of dengue virus | Emerging Infectious Diseases | No mosquito abundance data in low vs. high SES | Supplemental |
| 308 | Rochlin et al. | 2013 | Climate Change and Range Expansion of the Asian Tiger Mosquito (Aedes albopictus) in Northeastern USA: Implications for Public Health Practitioners | Plos One | Modeling paper | Supplemental |
| 309 | DeGroote and Sugumaran | 2012 | National and Regional Associations Between Human West Nile Virus Incidence and Demographic, Landscape, and Land Use Conditions in the Coterminous United States | Vector-borne and Zoonotic Diseases | No mosquito abundance data in low vs. high SES | WOS |
| 310 | Liu et al. | 2011 | Geographic incidence of human West Nile virus in northern Virginia, USA, in relation to incidence in birds and variations in urban environment | Science of the Total Environment | No mosquito abundance data in low vs. high SES | WOS |
| 311 | Rochlin et al. | 2011 | Predictive Mapping of Human Risk for West Nile Virus (WNV) Based on Environmental and Socioeconomic Factors | Plos One | No mosquito abundance data in low vs. high SES | WOS |
| 312 | Rey et al. | 2010 | Emergence of Dengue fever in America: Patterns, Processes, and prospects | Interciencia | Review | WOS |
| 313 | Vazquez-Prokopec et al. | 2010 | The Risk of West Nile Virus Infection Is Associated with Combined Sewer Overflow Streams in Urban Atlanta, Georgia, USA | Environmental Health Perspectives | No mosquito abundance data in low vs. high SES | WOS |
| 314 | Hayden et al. | 2010 | Microclimate and Human Factors in the Divergent Ecology of Aedes aegypti along the Arizona, U.S./Sonora, MX Border | Ecohealth | No mosquito abundance data in low vs. high SES | Supplemental |
| 315 | Liu and Weng | 2009 | An examination of the effect of landscape pattern, land surface temperature, and socioeconomic conditions on WNV dissemination in Chicago | Environmental Monitoring and Assessment | No mosquito abundance data in low vs. high SES | WOS |
| 316 | LaBeaud et al. | 2008 | Rapid GIS-based profiling of West Nile virus transmission: defining environmental factors associated with an urban-suburban outbreak in Northeast Ohio, USA | Geospatial Health | No mosquito abundance data in low vs. high SES | WOS |
| 317 | Ozdenerol et al. | 2008 | Locating suitable habitats for West Nile Virus-infected mosquitoes through association of environmental characteristics with infected mosquito locations: a case study in Shelby County, Tennessee | International Journal of Health Geographics | No mosquito abundance data in low vs. high SES | WOS |
| 318 | Ruiz et al. | 2007 | Association of West Nile virus illness and urban landscapes in Chicago and Detroit | International Journal of Health Geographics | Duplicate data (Ruiz et al. 2017) | Supplemental |
| 319 | Tackett et al. | 2006 | Relating West Nile Virus Case Fatality Rates to Demographic and Surveillance Variables | Public Health Reports | No mosquito abundance data in low vs. high SES | WOS |
| 320 | Rios et al. | 2006 | Demographic and spatial analysis of West Nile virus and St. Louis encephalitis in Houston, Texas | Journal of the American Mosquito Control Association | No mosquito abundance data in low vs. high SES | WOS |
| 321 | Brunkard et al. | 2004 | Dengue Fever Seroprevalence and Risk Factors, Texas–Mexico Border, 2004 | Emerging Infectious Diseases | No mosquito abundance data in low vs. high SES | WOS |
| 322 | Ruiz et al. | 2004 | Environmental and social determinants of human risk during a West Nile virus outbreak in the greater Chicago area, 2002 | International Journal of Health Geographics | No mosquito abundance data in low vs. high SES | Supplemental |
| 323 | Kutz et al. | 2003 | A geospatial study of the potential of two exotic species of mosquitoes to impact the epidemiology of West Nile virus in Maryland | Journal of the American Mosquito Control Association | No mosquito abundance data in low vs. high SES | WOS |
