## Appendix 2 for "Spatiotemporal patterns of urban mosquitoes are modulated by socioeconomic status and environmental traits in the United States"

*Ecological Applications*

Senay Yitbarek^1*^, Kelvin Chen^2^, Modeline Celestin^3^, and Matthew A. McCary^3^

^1^University of North Carolina at Chapel Hill, Department of Biology, Chapel Hill, NC 27599

^2^Rutgers University, Department of Ecology and Evolution, New Brunswick, NJ 08901

^3^Rice University, Department of BioSciences, Houston, TX 77005

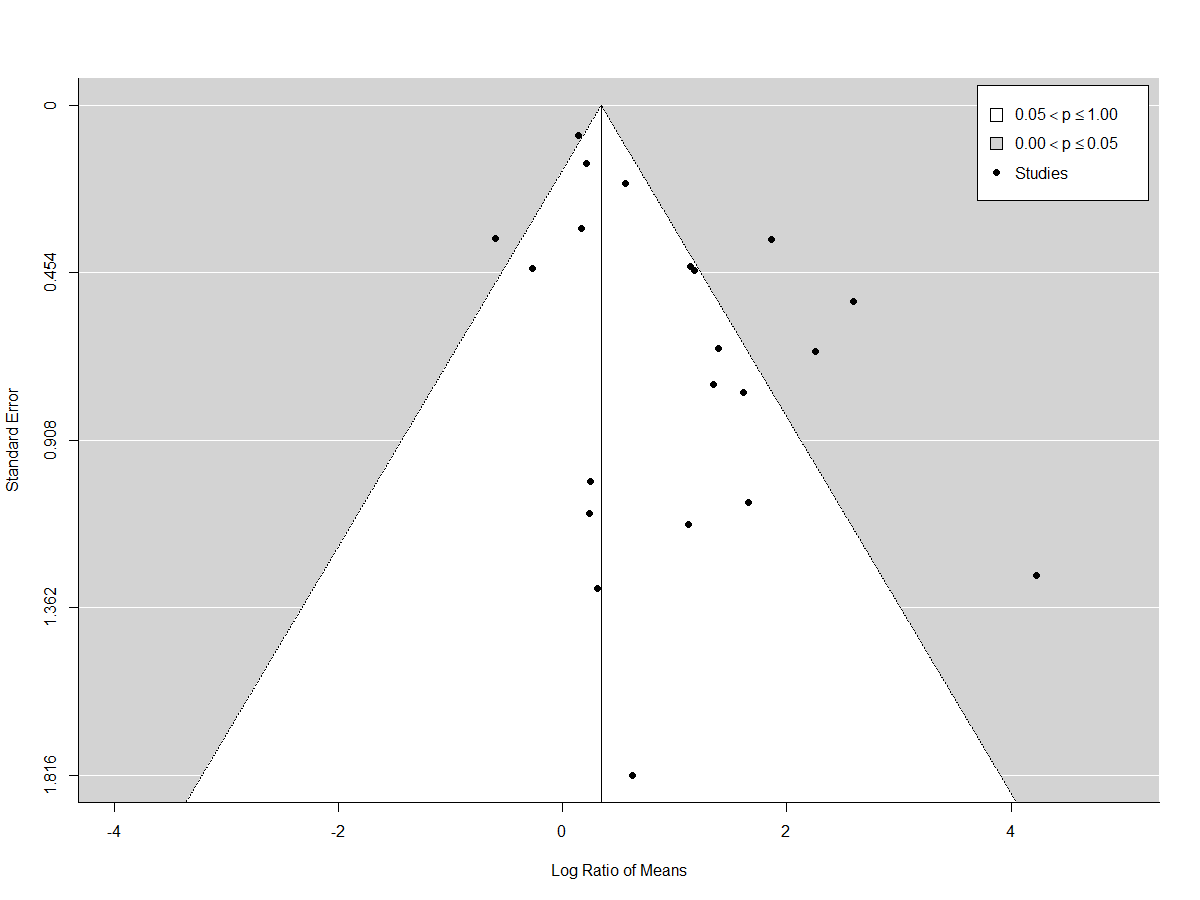

**Figure S1.** Trim-and-Fill funnel plot to illustrate potential publication bias for the influence of socioeconomic status (SES) on mosquito burden in urban environments. Here, the black dots show the spread of data from the meta-analysis. This Trim-and-Fill plot indicates a largely symmetrical pattern. The estimated number of missing studies on the left side is 9.

**Table S1.** The Rosenthal’s fail-safe number for the effects of SES on mosquito burden in urban environments. A *p*-value less than 0.05 suggests a rejection of the null hypothesis that a publication bias does exist in the data set.

| **Rosenthal's fail-safe number** | | ***p*-value** |
| --- | --- | --- |
| 441 | <0.001 | |
